# tRNA Analysis of eXpression (tRAX): A tool for integrating analysis of tRNAs, tRNA-derived small RNAs, and tRNA modifications

**DOI:** 10.1101/2022.07.02.498565

**Authors:** Andrew D. Holmes, Jonathan M. Howard, Patricia P. Chan, Todd M. Lowe

**Author notes:** These authors contributed equally to this work (Andrew D. Holmes, Jonathan M. Howard).

## Abstract

Recent high-throughput sequencing protocols have facilitated increased accuracy in measurements of transfer tRNAs (tRNAs) and tRNA-derived small RNAs (tDRs) from biological samples. However, commonly used RNA-seq analysis pipelines overlook special considerations given the unique features of tRNA metabolism. We present tRAX (tRNA Analysis of eXpression), a user-friendly analytic package for streamlined processing and graphic presentation of small-RNA sequencing data. Here, we apply it to both tRNAs and tDRs from mouse tissues to illustrate the extensive analysis and visualization features. Biologically compelling results demonstrate tRAX as an effective and accessible tool for in-depth characterization of tRNA and tDR transcriptomes.

## Background

While the canonical role of transfer RNAs (tRNAs) in mRNA translation has been well-studied [1–3], recent work has shown that the differential expression and modification of tRNAs, as well as their further processing into tRNA-derived small RNAs (tDRs; also known as tRFs) [4] can have important roles in cell homeostasis. For example, dysregulation of arginine (Arg) and glutamate (Glu) tRNAs has been shown to drive protein expression of key oncogenic factors in breast cancer models [5]. Additionally, modification of adenosine at position 58 (m^1^A_58_) can change in response to glucose levels as a means to regulate global mRNA translation during nutrient deprivation [6]. Other studies have found that increased generation of tDRs may correlate with cellular stress [7, 8], aging [9–11], and other biological processes and responses [12, 13]. tDRs may also affect gene expression through Argonaute-dependent silencing mechanisms [14, 15], reduced protein translation efficiency [8, 16], and increased mRNA degradation via competitive binding of proteins such as YBX1 [17]. These findings have motivated development of better methods to capture a global picture of tRNA and tDR dynamics.

High-throughput RNA sequencing has been a highly successful tool for studying expression of many types of genes. However, commercially available small RNA sequencing library preparation methods were primarily developed for microRNAs, and not designed to efficiently measure highly modified RNAs such as tRNAs. Some of the modifications commonly found on tRNAs, such as 1-methyladenosine (m^1^A) and 1-methylguanosine (m^1^G), may cause the reverse transcriptase to abort prematurely during cDNA synthesis [18], a required step for standard Illumina RNA sequencing library preparation. Depending on the protocol, a very large proportion of cDNAs will either be highly truncated or not amplified at all, leading to highly biased tRNA transcript representation. In some cases, modified bases in RNAs may appear as deletions and/or misincorporation of other bases in resulting cDNAs, possibly causing misalignment or non-alignment of sequencing reads [19]. Reverse transcription may also terminate due to inhibitory effects of highly stable secondary structure [20, 21]. Furthermore, library preparation protocols require RNA molecules to have a 3′ hydroxyl and 5′ phosphate moieties for ligation of sequencing adapters to transcripts. Mature, deacylated tRNAs have the necessary end chemistries [22], but pre-tRNAs have been found to carry incompatible 5′ triphosphates prior to RNase P cleavage of the leader sequence [23]. In addition, some tDRs may harbor 5′ hydroxyl, 3′ phosphates, or 2′3′-cyclic phosphates moieties via endonuclease fragmentation [24], resulting in an unquantified, potentially variable proportion of adapter ligation failures. Molecule end inaccessibility due to stable tRNA acceptor stems may further hinder adapter ligation [25]. Therefore, special measures are needed to achieve consistent, more accurate representation of tRNAs and tDRs in sequencing.

Aside from sequencing library preparation, unique features of tRNAs and tDRs complicate mapping and quantitation of sequencing reads. Standard small RNA-seq analytic methods were developed for microRNAs [26–29] and are not fully applicable to tRNAs. As one of the largest gene families in most genomes, a complex assortment of tRNA genes must encode at least thirty different types of tRNAs, accurately decode at least sixty one codons, and convey at least twenty different amino acids to the ribosome. While small genomes of some bacteria and archaea may have as few as about 30 unique tRNA genes [30–32], tRNA gene copy and complexity increases in multicellular organisms, with the human genome having 500+ tRNA genes [30–32]. For example, in terms of gene redundancy, seventy human tRNAs (GRCh38) have at least one other identical copy in the genome [30]. Vertebrates also tend to have many tRNA isodecoders (tRNAs with the same anticodon but slightly different sequences in the gene body), with sequences containing only one or a few nucleotide differences. tDRs that are derived from multi-copy tRNA genes, as well as those processed from a region identical across multiple isodecoders will have an ambiguous origin. Therefore, it is often impossible for sequencing reads to be uniquely aligned to a specific tRNA locus. Analysis tools that only consider uniquely mapped sequencing reads for abundance estimation will miss a large portion of those belonging to tRNAs or tDRs, depending on the genome. Algorithms that assume small RNAs not containing introns can also result in incomplete or incorrect read mapping, as some archaeal and eukaryotic tRNA genes carry introns. Furthermore, all eukaryotic tRNAs require post-transcriptional addition of 3′ CCA tail during maturation to facilitate aminoacylation [33–35], in contrast to bacterial and archaeal tRNAs that may or may not have a genomically encoded 3′-terminal CCA tail. Sequencing reads for mature tRNAs or tDRs derived from the 3′ end of tRNAs may, therefore, not be able to fully align correctly to the reference genome. Thus, an analysis tool specially designed for tRNA sequencing data is needed to produce the most accurate results.

Recently published tRNA-specific sequencing methods have introduced solutions to address these difficulties. For example, ARM-seq (AlkB-facilitated RNA methylation sequencing) [36] and DM-tRNA-seq (Demethylase-thermostable group II intron RT tRNA sequencing) [37] use RNA pre-treatments with wildtype or mutationally-enhanced versions of *E. coli* AlkB to remove commonly methylated tRNA residues, facilitating read-through of reverse transcriptase. In addition, the DM-tRNA-seq library preparation protocol utilizes a highly processive thermostable group II intron reverse transcriptase (TGIRT) and relies on ligation-independent, end-to-end template switching during adapter attachment. While TGIRT extends through most types of RNA modification, it also produces misincorporations at these modifications that can be detected and utilized to predict modifications in sequencing data [38, 39]. More recently, the OTTR-seq library preparation protocol has married the aforementioned protocols, allowing sequencing of both tDRs and mature tRNAs from the same sequencing pool, with the added advantage of demarcating the two subtypes [40]. Special software tools including MINTMap [41], SPORTS [42], and tDRmapper [43] have been developed, yet they focus on tDRs to the exclusion of mature tRNAs. To address the challenges of tRNA sequencing analysis, we have developed tRAX: tRNA Analysis of eXpression. In addition to detecting expression of a wide variety of annotated small RNAs such as microRNAs, this software package provides in-depth tRNA and tDR abundance quantification and differential expression analysis across any number of experimental samples or replicates. It also detects misincorporations within DM-tRNA-seq (or any other highly processive RT enzyme) data for predicting RNA modifications in tRNAs. We demonstrate and discuss new insights made possible from tRAX analyses, leveraging over fifty types of visualizations to profile and compare newly collected data across multiple mouse tissues. With the comprehensive features and detailed user guide, tRAX can be an invaluable tool to enable all tRNA and tDR researchers to comprehensively explore their own high-throughput small RNA sequencing data sets.

## Results

### Overview of tRAX analysis of tRNA-sequencing data

tRAX is a software package built for in-depth analyses of tRNA-derived small RNAs (tDRs), mature tRNAs, and inference of RNA modifications from high-throughput small RNA sequencing data. At its core, the tRAX workflow (Additional file 1: Figure S1) incorporates a complete set of RNA sequencing “best practices” functionality including adapter trimming, read alignment to reference, transcript abundance estimation, and differential expression analysis [44] for all annotated small RNAs. Importantly, it integrates special features to support the characteristics of tRNAs and tDRs. For example, sequencing reads are aligned to a custom-built reference database including all tRNA loci in the reference genome (matching all pre-tRNAs), as well as fully-processed mature tRNA transcripts with 3′ CCAs tails. Sequencing read mappings are prioritized to mature tRNA transcripts, followed by reads aligning to genomic loci that cover all precursor tRNAs and other ncRNAs such as microRNAs, rRNAs, snoRNAs, snRNAs. Unlike standard RNA-seq read counting tools that only consider or recommend uniquely mapped reads for quantitation [45, 46], tRAX allows reads to be mapped to multiple transcripts and gene loci, which is necessary to include the major contributions of identical or near-identical tRNA genes. tRAX innovates the classification of tRNA read coverage in four categories: transcript-specific, isodecoder-specific (equal matches to multiple different tRNAs with the same anticodon), isotype-specific (equal matches to tRNAs decoding the same amino acid), and those spanning multiple isotypes. In order to also report the abundance of major tRNA fragment types, tRAX computes separate read counts for those that align at the tRNA 5′ end, 3′ end, or “other” regions not anchored to either end. These data are used to compute differential expression across samples for both full-length tRNA transcripts and tDRs, enabling scientists to recognize changes in fine detail. tRAX also measures base frequencies of reads at each position of each tRNA to determine reverse transcriptase misincorporation rates. Used in combination with enzymes that are highly processive such as TGIRT, Superscript IV, and others [37, 38, 40, 47], these analyses show characteristic patterns for numerous RNA modifications that are integral to tRNA function, stability, and regulation [48–50].

tRAX generates data in multiple formats at each stage of the analysis (Additional file 2: Table S1). First, a quality assessment (QA) report is provided for an estimation of the quality of sequencing data used for analysis (for example, see Additional file 3). It contains statistics we commonly use to reveal problems in library preparation or sequencing, with each metric falling into a “Passed”, “Warning”, or “Failed” value range (see Methods). In addition to profiling tRNAs and tDRs, tRAX estimates the relative abundance of other annotated small RNAs, summarized in histograms and scatter plots to give better context to tRNA abundance. Normalized read counts and differential expression analysis results including log_2_ fold change and adjusted P-value of annotated small RNAs are provided in tab-delimited files. To visually explore and compare changes in transcripts among samples, tRAX generates read coverage plots for each unique tRNA isodecoder as well as data tracks showing transcript abundance for each tRNA gene, directly viewable in the UCSC Genome Browser [51, 52]. Moreover, principal component analysis and volcano plots are available to further profile changes. To screen for RNA modifications or editing events, potential nucleotide misincorporations are presented in a series of graphical dot plots and heatmaps to uncover characteristic patterns by tRNA position, isotype, isodecoder, and sample. Taken together, tRAX provides unrivaled flexibility to gain novel insights into tRNA expression, processing, and regulation by creating a multitude of germane statistical analyses, sample comparisons, and tRNA-focused graphical visualizations.

### Analysis of tRNA expression in mouse tissues as an illustration of tRAX functions

To highlight the functionality and insights provided by tRAX, we performed tRNA sequencing of three contrasting tissue types from wildtype mice, including brain, heart, and liver. Previously developed tRNA-specific sequencing methodologies ARM-seq [36] and DM-tRNA-seq [37] were employed to capture the expression of tDRs and mature tRNAs, respectively. Both protocols include the use of AlkB to remove specific RNA base methylations that can impair efficient reverse transcription [18] (Fig. 1A). To fully leverage the statistical power of biological replicates for data analysis, tissue samples were obtained from multiple mice. After performing deep sequencing of these samples, we used tRAX to assess tissue-specific differences in tRNA and tDR expression, as well as to identify likely RNA modifications.

**Fig. 1.**
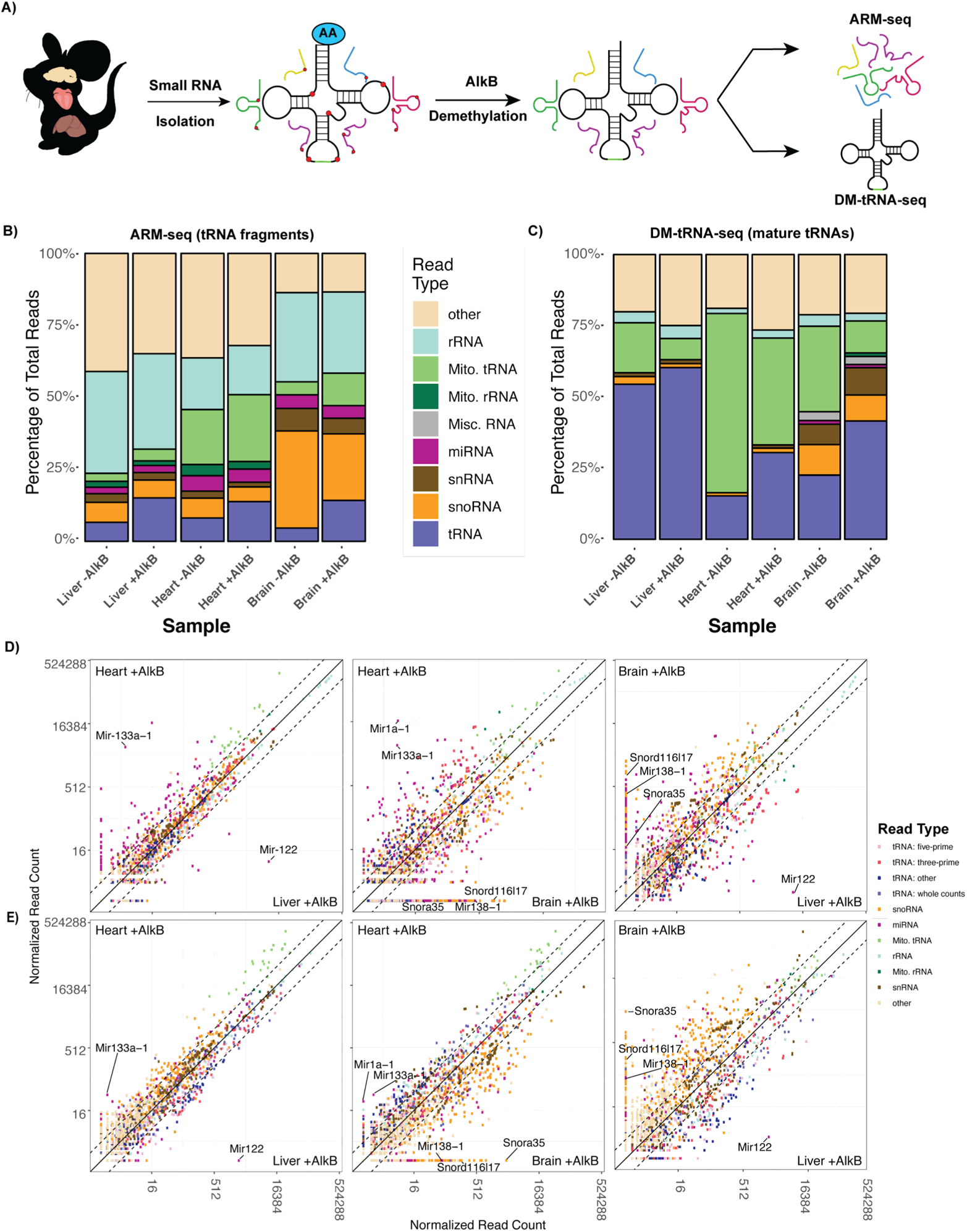
tRAX pipeline workflow and general analysis of ARM-seq and DM-tRNA-seq data. **A** Graphical workflow for mouse tissue-specific tRNA sequencing. Mouse heart, liver, and brain samples were harvested for total RNA isolation. Small RNAs were then isolated and treated under control (-AlkB) and demethylation (+AlkB) conditions (methylations shown as *red circles*). These RNAs were then subjected to ARM-seq and DM-tRNA-seq sequencing library preparations (see *Methods*). **B** Sequencing read distribution across RNA categories for −AlkB and +AlkB samples from mouse liver, heart, and brain ARM-seq sequencing libraries. **C** Sequencing read distribution across RNA categories for −AlkB and +AlkB samples from mouse liver, heart, and brain from DM-tRNA-seq sequencing libraries. **D,E** Normalized read count scatter plots comparing differences in abundance of RNAs detected between tissues for (**D)** ARM-seq and (**E)** DM-tRNA-seq. Example tissue-enriched RNAs are labeled to highlight abundance differences with respect to sequencing protocol.

Beginning with summaries generated by tRAX, we noted clear differences between the ARM-seq and DM-tRNA-seq experiments in terms of sequencing read distributions across mouse tissues for various types of non-coding RNAs, viewable by relative proportions (Fig. 1B-C) or by absolute read counts (Additional file1: Figures S2, S3). ARM-seq libraries (Fig. 1B) included many more non-coding RNA types than DM-tRNA-seq libraries (Fig. 1C) which we estimate is mostly due to differences in reverse transcriptases and protocol bias for or against full-length tRNAs. Mature tRNAs make up the largest proportion of small RNAs in cells [53], thus DM-tRNA-seq’s TGIRT enzyme captures this relative abundance faithfully (Fig. 1C). By contrast, ARM-seq’s less processive RT enzyme disfavors highly modified mature tRNAs, producing a more balanced representation among all small RNA types (Fig. 1B). Another difference is apparent in the increased tRNA read portion for samples with AlkB treatment (+AlkB) versus those without (-AlkB), for both ARM-seq and DM-tRNA-seq. This further confirms that AlkB demethylation facilitates the read-through of modified tRNAs and tDRs [36]. Visualizations of read length distributions for both forms of sequencing show that ARM-seq indeed recovers short and medium length tRNA fragments, along with many other RNA types and sizes (Additional file 1: Figures S4) while DM-tRNA-seq preferentially profiles mature full-length tRNAs (Additional file 1: Figures S5). The sequencing read length distributions for both tRNA-derived and other RNAs are remarkably different and reproducible across the three tissues tested.

Because tRNA transcript levels have only recently been observed effectively with new tRNA-seq methods, it is important to place them in context of previously studied non-coding RNAs. tRAX provides normalized read count scatterplots (Fig. 1D-E) and volcano plots (Additional file1: Figures S6, S7) for rapid identification of sample type-specific patterns for all annotated small RNAs. These plots are complex, but large shifts and outliers are readily apparent. In our three-tissue comparison, one of the more striking results is visible in ARM-seq plots (Fig. 1D) but most evident in DM-tRNA-seq plots (Fig. 1E), both showing a much higher abundance of numerous snoRNAs in brain tissue relative to heart and liver. Among these, previously studied brain-specific Snora35/MBI-36 [54] (human ortholog: HBI-36) exhibited a log_2_ fold change (LFC) value above 12.0 when comparing brain to liver or heart tissue (Fig.1E, Additional file1: Figure S7, and Additional file 4: Table S2). Similarly, snoRNAs derived from the Snord116 gene cluster [55] were measured to have LFCs above 8.0 in brain relative to the other tissues (Fig. 1E, Additional file 1: Figure S7, and Additional file 4: Table S2). We also observed consistency with previously reported tissue-specific miRNAs in both ARM-seq and DM-tRNA-seq results, including brain-specific miR-138 [56] (LFCs > 7.3), liver-enriched miR-122 [57] (LFCs > 6.9), and heart- (and generally muscle-) enriched miR-133a [58] (LFCs > 6.0) (Fig. 1D, Additional file 1: Figure S6, S7, and Additional file 4: Table S3, S4). These “positive control” examples demonstrate tRAX’s general ability to detect tissue-specific small RNA transcripts within our samples, as measured by either tRNA-seq method.

### tRAX facilitates discovery of tissue-specific tRNA-derived small RNAs

To study tDR abundance in mouse tissues utilizing ARM-seq, we started by examining the sequencing read distribution (Additional file 1: Figure S8) and per-base coverage categorized by tRNA isotypes (Additional file 1: Figure S9) generated by tRAX. While tDRs in AlkB-treated (+AlkB) samples had a net average of 2.5 fold more sequencing reads than those in the untreated (-AlkB) samples, some tRNA isotypes had more dramatic gains than others (Additional file 1: Figure S8). tDRs derived from tRNA^Phe^ had almost an average of 10 times more sequencing reads in +AlkB samples, and those derived from tRNA^Arg^ had over 6.7 times as many, indicating that these tDRs harbor many more AlkB-sensitive RNA modifications that impede reverse transcription. This is further visualized in the per-base coverage plot where the sharp dropoff near the 3’ end of the tRNA in the −AlkB samples, commonly found with m^1^A_58_ modification [59], was replaced by a smoother curve gradually decreasing into the anticodon loop region in the +AlkB samples (Additional file 1: Figure S9).

Within these aggregate read plots, the uneven sequencing read coverage along the full length tRNA region raises the question of which specific tDRs differ among the three tissue types. To address this, tRAX produces isotype-labelled transcript scatterplots subsetting sequencing reads derived from the 5′ region, 3′ region, internal region, and full-length tRNAs for all pairwise tissue comparisons (Additional file 1: Figure S10 – S12). Upon examination of these plots, we noted tissue biases varied markedly depending on the type of tDR: 5’ tDRs are relatively more abundant in brain than heart (Fig 2A, 2D), yet the opposite tissue bias was observed for 3’ tDRs in heart versus liver (Fig. 2B, 2E), and those derived from internal tRNA regions in heart versus brain (Fig. 2C, 2F). Because tRAX creates detailed read coverage plots for every tRNA for every sample, we are able to examine the precise start and end of any tDR of interest from the scatterplot. Looking at tRNA-Val-AAC-4 as an example (red circle highlighted in Fig. 2A), the 5’ tDR derived from this valine tRNA has sequencing reads in the brain that are almost entirely absent in heart (Fig. 2D) and liver (Additional file 1: Figure S13A). Reads in these coverage plots are classified by how specifically the RNA-seq reads map to one or more tRNA transcripts: purple indicates uniquely mapping reads (no ambiguity); cyan indicates reads mapping to multiple tRNAs that all have the same anticodon; green represents multiple mapping to tRNAs of the same isotype but different anticodons, and salmon indicates non-specific reads that map across tRNAs from multiple isotypes. For tRNA-seq analysis, knowing the ambiguity of tRNA reads is essential for ascertaining the source(s) of any given tDR of interest. The uniqueness of the 5’ tDR mapping to tRNA-Val-AAC-4 (Fig. 2D, purple coverage in Brain +AlkB) is due to a U_6_:A_67_ pair in the acceptor stem (Additional file 1: Figure S13C). The coverage plot for this tDR resembles a “tRNA half” from the 5′ end (position 1) to the anticodon (position 36) (Additional file 1: Figure S13B) for brain samples. While the function of this specific tDR is still unknown, previous studies reported correlated changes in abundance of tRNA^Val(AAC)^ 5′ halves with onset of cellular stress [60] and within cancer models [61], as well as potential for regulatory roles in gene expression of aging mammalian brain [10].

**Fig. 2.**
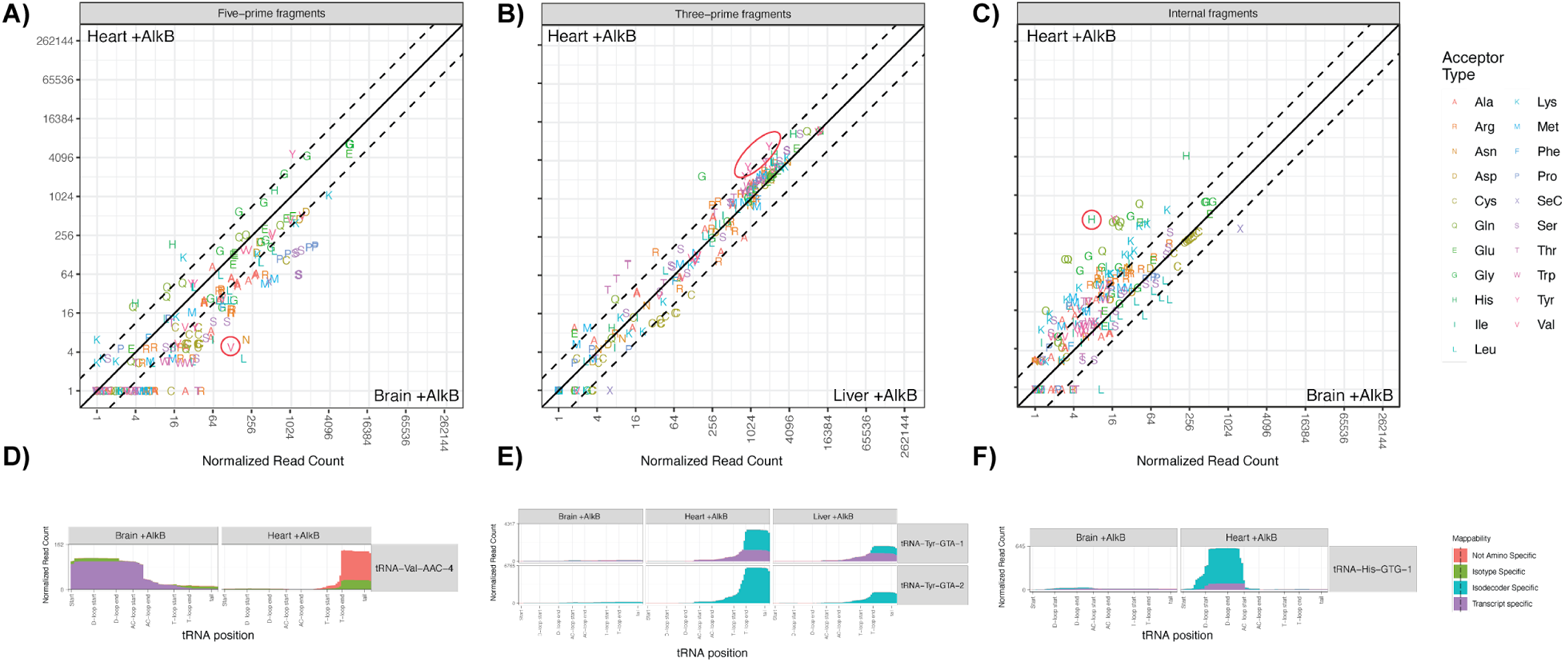
Tissue-enriched tDR expression profiles in mouse heart, brain, and liver. **A,D** Brain-enriched 5’-derived tDR from tRNA-Val-AAC-4. **A** Scatter plot of normalized read count comparison between heart and brain ARM-seq samples, with tRNA-Val-AAC-4 highlighted (red circle). **D** Per-base read coverage of tRNA-Val-ACC-4 in mouse brain and heart. **B,E** Heart-enriched 3’-derived tDR from tRNA-Tyr-GTA isodecoders. **B** Scatter plot of normalized read count comparison between heart and liver ARM-seq samples, with tRNA-Tyr-GTA isodeocoders highlighted (red circle). **E** Per-base read coverage of tRNA-Tyr-GTA-1 and -2 in mouse brain, heart and liver. **C,F** Heart-enriched internally-derived tDR from tRNA-His-GTG-1. **C** Scatter plot of normalized read count comparison between heart and brain ARM-seq samples, with tRNA-His-GTG-1 highlighted (red circle). **F** Per-base read coverage of tRNA-His-GTG-1 and -2 in mouse brain and heart.

In contrast, tRNA-Tyr-GTA-1 and tRNA-Tyr-GTA-2 tDRs were lowly expressed in brain samples (Fig. 2E, left panel) but abundant in heart and liver (red circle highlight in Fig. 2B, 2E, and Additional file 1: Figure S11, S12). Because these two isodecoders share the same sequence in the 3’ end except a one-base difference at the T-loop (position 59), both uniquely (purple) and non-uniquely aligned (cyan) sequencing reads for 3’ tDRs are observed. Similar to those found in previous human and mouse studies [36, 62–65], these tDRs have different 5′ start sites along the T-loop of tRNA^Tyr(GTA)^ but end consistently at the 3′ CCA tail (Additional file 1: Figure S14A). Further studies will be needed to understand the significance of the tDR sequence variants in terms of function and regulation. Likewise, tDRs derived from the “internal” region of mature tRNAs have not been commonly found and studied [66]. Interestingly, we observed heart-specific tDRs from tRNA-His-GTG-1 through the internal fragment subset of the tRAX results (Fig. 2C and 2F). While these “internal” tDRs also have different 5′ start sites, the position variability may suggest distinct processing and regulatory mechanisms (Additional file 1: Figure S14B). Previous research suggests that human 5′ halves, including the one derived from tRNA^His(GUG)^, may be angiogenin-responsive [67]. Investigating the potential relationship between the “internal” tDRs and angiogenin could be one follow-up study prompted by these tRAX results.

### End chemistry analysis of tDRs using PNK enzymatic treatments

Current commercial and specialized sequencing library preparation methods depend heavily on particular nucleic acid end chemistries to optimize target molecule inclusion in the final sequencing pool. While many cellular RNA species contain 5′ phosphate and 3′ hydroxyl moieties that are needed for commonly used library preparation kits, there exist many other RNA molecules with different end chemistries due to variation in cellular processing or enzymatic treatments prior to the library preparation. 5′ tDRs that are generated by angiogenin (ANG) cleavage, leaving 2′,3′-cyclic phosphates and 5′ hydroxyl moieties [68] may result in a substantial proportion of incompatible ends for standard ARM-seq library preparation. To assess this issue, we included an additional ARM-seq library preparation on mouse liver RNA samples that incorporated a T4 polynucleotide kinase (T4 PNK) treatment in the absence of ATP and at low pH [69] (Fig. 3). These specific conditions promote the 3’ phosphatase/2’,3’-cyclic phosphatase activity of T4 PNK to restore the required end chemistry (see Methods). tRAX enabled us to compare samples treated with both AlkB and T4PNK (+AlkB/+PNK), those only treated with AlkB (+AlkB), and those without any treatment (-AlkB). When reviewing the overall per-base coverage for all tRNAs, we noticed that +AlkB/+PNK samples had a pronounced increase of sequencing reads in the 5′ half region that was at very low level in the samples without PNK treatment (Fig. 3A). This general increase in 5′ coverage of tRNAs also coincides with almost doubling the proportion of reads mapping to tRNAs (Additional file 1: Figure S15), indicating that the T4 PNK treatment effectively facilitates the inclusion of these 5’ tDR subtypes which would be otherwise missed [24, 70]. We also note that the increase in 5’ tDRs dilutes the abundance of many 3’ tDRs, thus increased read coverage may be needed to measure low-abundance 3’ tDRs.

**Fig. 3.**
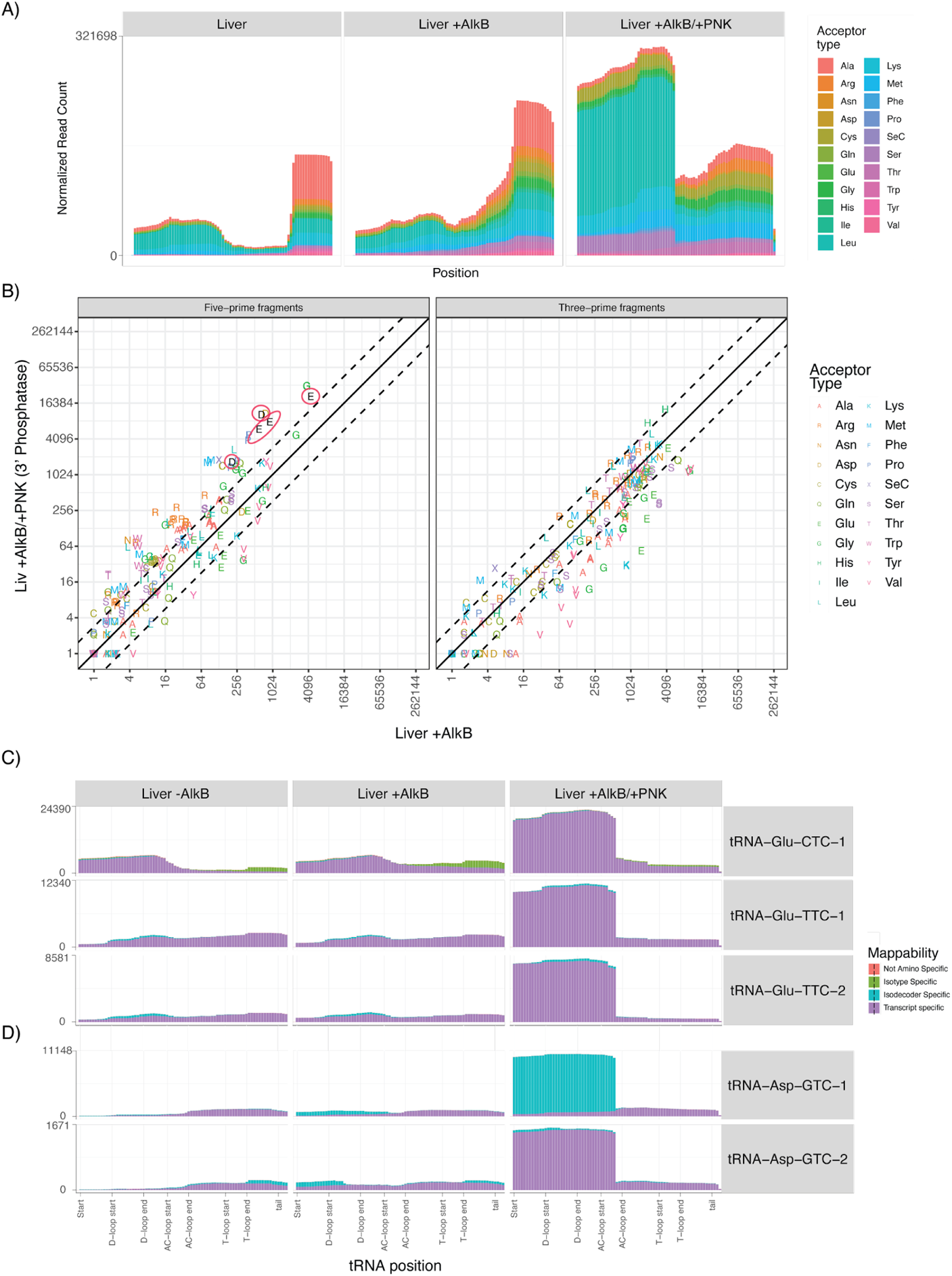
Treatment of T4 polynucleotide kinase (PNK) reveals a subset of tDRs in mouse liver. **A** Combined read coverage across all tRNAs for −AlkB, +AlkB, and +AlkB/+T4PNK-treated samples from mouse liver ARM-seq experiments. **B** Scatter plot of normalized read count comparison between +AlkB and +AlkB/+T4PNK treated samples for 5’ and 3’ tRNA fragment types. **C,D** Per-base read coverage of (**C**) tRNA-Glu-CTC/TTC and (**D**) tRNA-Asp-GTC isodecoders for −AlkB, +AlkB, and +AlkB/+T4PNK-treated samples.

To determine whether this increase in 5′ coverage was common to all tRNAs or specific ones, we turned to the abundance comparison between +AlkB/+PNK and +AlkB only samples, separated into 5′ or 3′ fragments (Fig. 3B). Generally, with phosphatase treatment using T4 PNK, an increase in 5′-derived tDRs was observed for most of the tRNAs. Some of the highly abundant 5′ tDRs include those derived from tRNA^Glu^ and tRNA^Asp^ (Fig. 3B, right panel; Fig 3C-D). Looking at these examples more closely, we noticed that both sets of 5′ tDRs terminate at position 36 (Additional file 1: Figure S16) between the dinucleotide “CA” (Additional file 1: Figure S17), which is the preferred sequence of endoribonuclease angiogenin [68, 71]. This further supports the likely angiogenin-driven origin of these tDRs, and illustrates the utility of T4 PNK treatment to enable their sequencing. In contrast, we see the 3′ tDR enrichment bias with +AlkB only treatment (Fig. 3B, right panel). These findings drive home that ARM-seq and other common small RNA sequencing methods can be heavily biased toward different fragment types. These data illustrate that sequencing output can be heavily dependent on enzymatic treatments of sample RNA and library preparations; thus, multiple methods may be needed to gain a more complete view of the small RNA transcriptome. Overall, this example shows how effective tRAX is for evaluating the breadth and depth of differences between sample preparation methods.

### tRAX highlights expression of tissue-specific mature tRNAs

In metazoans, tissue-specific expression of protein coding genes is an essential foundation for enabling different cell types to take on specialized functions. A landmark study found tRNA-Arg-TCT-4 (one out of five isodecoders producing Arg-UCU tRNAs) was exclusively abundant in mouse brain [72]. Unexpectedly, mutations in this tRNA increased seizure threshold and altered synaptic transmission when combined with impared ribosome rescue factor GTPBP2 [72, 73]. To investigate the existence of tissue-biased tRNAs in mice, we used tRAX to examine the expression of mature tRNAs in brain, heart, and liver with DM-tRNA-seq. The sole standout brain-enriched tRNA was the previously noted tRNA-Arg-TCT-4, with over six-fold greater abundance than the other tissues (Fig. 4A). The individual transcript read coverage plots further documented the stark difference in expression among the tRNA-Arg-TCT isodecoders (Fig. 4C). Although tRNA-Arg-TCT-4 was not as abundant in brain as most of the other isodecoders, it was the only one that exhibited substantial expression in brain relative to other tissue samples. This agreement with the best-documented case of tissue-specific tRNA expression may be considered a positive control supporting both the sequencing method and tRAX analytics.

**Fig. 4.**
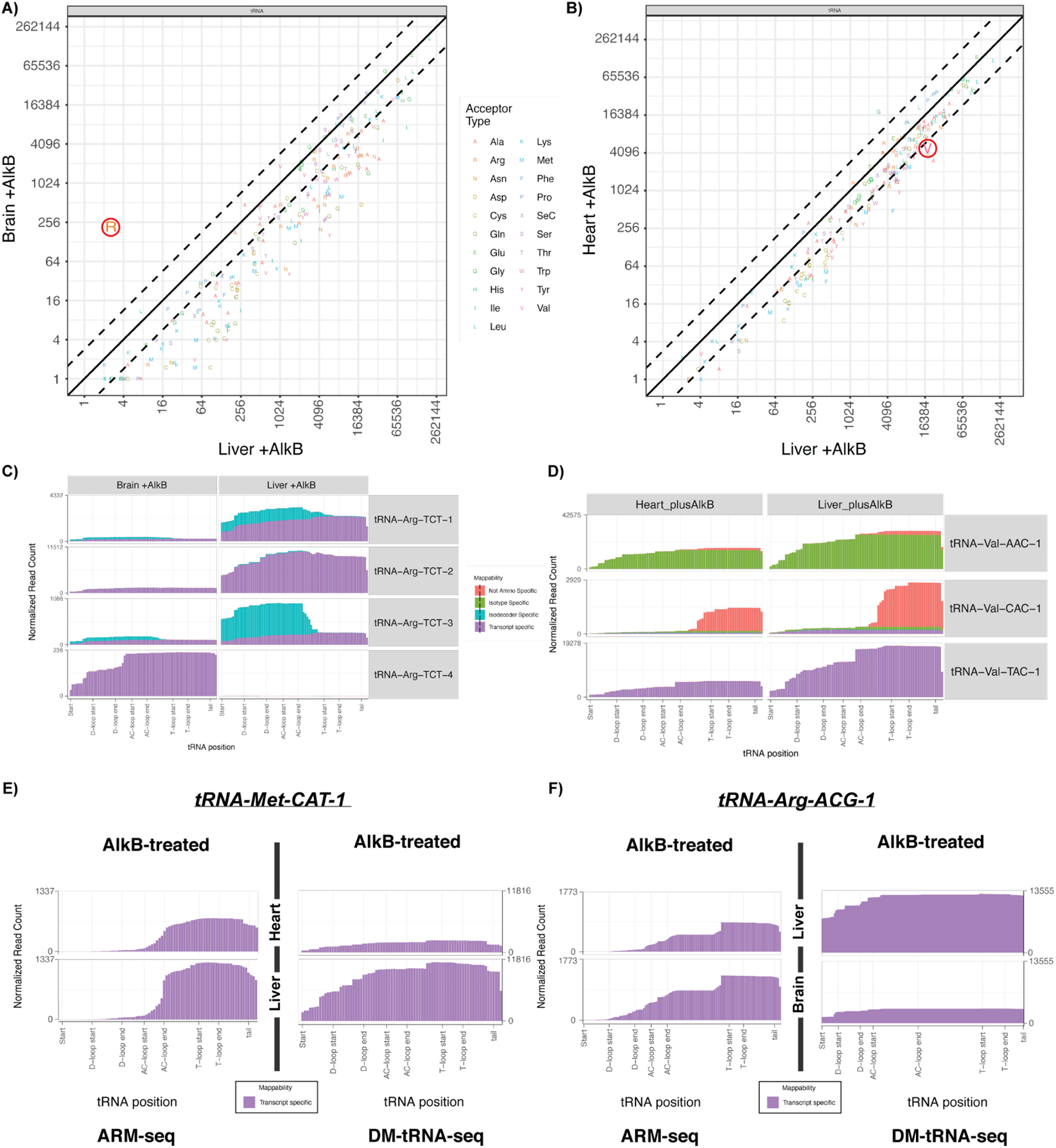
Tissue-enriched mature tRNA expression profiles. **A,C** Brain-enriched tRNA-Arg-TCT-4. **A** Scatter plot of normalized read count comparison between brain and liver DM-tRNA-seq samples, with tRNA-Arg-TCT-4 highlighted (red circle). **C** Per-base read coverage of tRNA-Arg-TCT isodecoders in mouse brain, heart, and liver. **B,D** Liver-enriched tRNA-Val-TAC-1. **B** Scatter plot of normalized read count comparison between heart and liver DM-tRNA-seq samples, with tRNA-Val-TAC-1 highlighted (red circle). **D** Per-base read coverage of tRNA-Val-TAC-1 in mouse brain, heart, and liver. **E** Comparison of AlkB-treated ARM-seq and DM-tRNA-seq read coverage for tRNA-Met-CAT-1. **F** Comparison of AlkB-treated ARM-seq and DM-tRNA-seq read coverage for tRNA-Arg-ACG-1.

Additional examination of the overall abundance comparison across the three tissue types reveal that many tRNAs were enriched in liver (Fig. 4B). For example, we observed that tRNA-Val-TAC-1 was twice more abundant in liver than heart (Fig. 4D). Interestingly, tDRs derived from the human tRNA-Val-TAC isodecoders were found to be potential plasma biomarkers associated with liver cancer [74]. To examine if this might also apply to our mouse liver samples, we compared the read coverage between ARM-seq and DM-tRNA-seq data for liver and heart and found an inverse relationship, suggesting that the majority of tRNA-Val-TAC-1 in the liver was in the mature form (Additional file 1: Figure S18). Overall, the above two specific examples, plus the variable relative amounts of many other individual tRNAs (Fig. 4A-B) substantiate that tissue-enriched tRNAs are more common than previously anticipated, and imply a larger unexplored role for tissue-biased tRNAs in regulating translation and other cellular processes.

With tRAX analysis, we also have an opportunity to evaluate the relative abundance levels of mature tRNAs to their tDRs [75]. Overall, we observed complex, varying relationships between the two. For example, when comparing ARM-seq to DM-tRNA-seq data for tRNA-Met-CAT-1, we observed that the general ratio of tDRs (ARM-seq) derived from the 3’ region of this tRNA between heart and liver tends to follow the ratio of mature (DM-tRNA-seq) tRNA reads; that is, more mature tRNA transcripts produce more tDRs (Fig. 4E). However, the abundance ratio of mature tRNA-Arg-ACG-1 between liver and brain was inversely correlated to its tDR counterpart (Fig. 4F). Therefore, our preliminary analyses indicate that tissue-biased mature tRNAs and their corresponding tDRs have their own unique expression, processing, and stability profiles, suggesting regulatory roles that vary depending on tissue context. tRAX is designed to illuminate these complex relationships and power future characterization studies.

### tRAX detects tRNA modifications via quantification of RT misincorporation and terminations

Human cytosolic tRNAs have been shown to contain anywhere from 3 to 17 modifications [12]. While many modifications are not visible in sequencing data, some of them cause partial misincorporation and/or deletions of nucleosides during the reverse transcription step of cDNA library preparation. For example, adenosine-to-inosine (A-I) RNA editing leads to a guanosine in cDNA instead of the adenosine in the original genomic sequence. The mismatches noted at read alignments can be used to detect the modification state of tRNAs at single-base resolution. tRAX employs this idea and allows us to discover and assess RNA modifications from any tRNA-seq method that employs highly processive RT enzymes. In this study using DM-tRNA-seq, we started by exploring the misincorporations detected at tRNA position 9, which is known to harbor m^1^A or m^1^G in some tRNAs [59, 76–78]. By comparing between AlkB-treated (+AlkB) and untreated (-AlkB) samples from mouse brain, heart, and liver, we observed a multi-fold decrease in misincorporation percentage in +AlkB samples (Fig. 5A), demonstrating the effectiveness of AlkB demethylase treatment and confirmation of methylation at this position for a subset of tRNAs. Evaluating the same data at the tRNA isotype level showed that the percentage of misincorporations varied remarkably, with tRNA^Arg^, tRNA^Asp^, and tRNA^Ile^ having a relatively larger frequency of misincorporation (thus inferred modification) at position 9 (Fig. 5B), as shown in previous studies [59, 77, 78]. When we expanded the view to all positions in all mature tRNAs, we found that positions 9, 26, 34, 37, and 58 had the widest spread of misincorporations among the different tRNA isotypes (Fig. 5C). These potentially represent some commonly known tRNA modifications, such as m^1^G_9_, m^2^ G, and m^1^A [79]. Whereas, the scattered number of isotypes at some positions like 47 and 49 reflect the modifications such as m^3^C_47_ and m^5^C_49_ that were previously found only in some specific tRNAs [59, 78, 79].

**Fig. 5.**
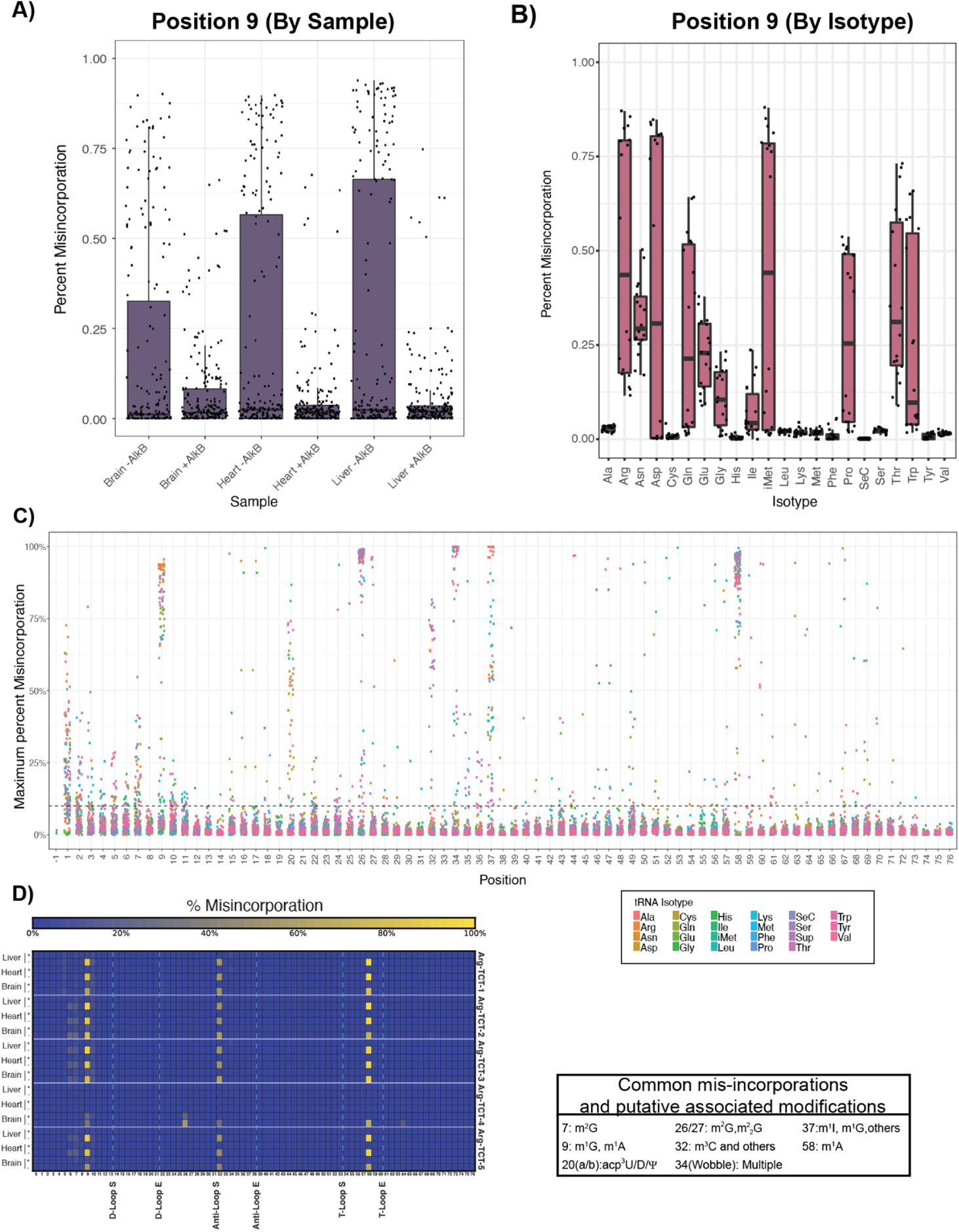
Misincorporations detected in DM-tRNA-seq data reveal putative modification states in tRNAs. **A** Percentage of sequencing reads with nucleotide misincorporations at position 9 in all expressed tRNA transcripts for mouse liver, heart, and brain AlkB-treated (+AlkB) and untreated (-AlkB) samples. **B** Percentage of sequencing reads with nucleotide misincorporations at position 9 categorized by tRNA isotype for all studied tissues (mouse heart, liver, and brain) in −AlkB samples. **C** Percentage of misincorporation at each tRNA position for all studied samples. Each dot represents an isodecoder categorized by tRNA isotype. **D** Heatmap of misincorporation percentages for tRNA-Arg-TCT isodecoders in +/-AlkB samples of mouse liver, heart, and brain.

When comparing misincorporation levels in our samples across the tRNA-Arg-TCT isodecoders, it is obvious that all but the brain-enriched tRNA-Arg-TCT-4 had comparable misincorporations in terms of percentages and positions (Fig. 5D), suggesting that these isodecoders may have the same RNA modifications in mouse brain, liver, and heart. However, the detected amount of misincorporations at positions 9 and 32 of tRNA-Arg-TCT-4 diminished while there was a gain of misincorporation at position 26. As mentioned before, m^2^ G has been found at position 26 of multiple tRNAs [81, 82]. The misincorporation evidence for m^2^ G in tRNA-Arg-TCT-4 (a modification absent in the other Arg-UCU isodecoders) may suggest a link between the modification and the tissue-specific role of the gene. This finding shows the capability of tRAX for identifying novel modification sites in tRNAs, and clear differences in modifications between isodecoders.

In conjunction with misincorportations, the tRAX results allow us to determine reverse transcription (RT) terminations at each position across all sequenced tRNAs. While not all RT terminations represent RNA modifications or degradation products, over-representation of sequencing read ends that do not terminate at the reverse transcription end of a mature tRNA (5’ read end at tRNA position 1) suggests potential early read-through failure due to modifications. One example is tRNA^Ala^ which harbors an N^1^-methylinosine (m^1^I) modification at position 37 [80]. Analysis of the 5′ ends in our mouse DM-tRNA-seq data revealed a large increase in RT terminations at position 38 (Fig. 6A), which appeared to be sensitive to AlkB demethylation. Notably, when comparing the detected level of 5′ ends with the amount of misincorporations at each position of an exemplary tRNA^Ala^ (tRNA-Ala-AGC-4), we noticed RT terminations built-up at position 38 with high level of misincorporations at the adjacent position 37 (Fig. 6B). These terminations were eliminated by AlkB treatment while the levels of misincorporations increased slightly with demethylation. This is possibly because methylated inosine leads to misincorporation of multiple bases (including the correct adenosine base) by TGIRT, versus misincorporation of guanosine due to the remaining inosine edit after demethylation [78]. These data show that not only can TGIRT enzyme process through multiple types of tRNA modifications, but that misincorporations and RT terminations detected in tRAX results can be used together to ascribe specific sequencing aberrations to specific tRNA modifications. Additional methods such as liquid chromatography tandem mass spectrometry (LC-MS/MS) can be further applied to verify the newly detected modifications.

**Fig. 6.**
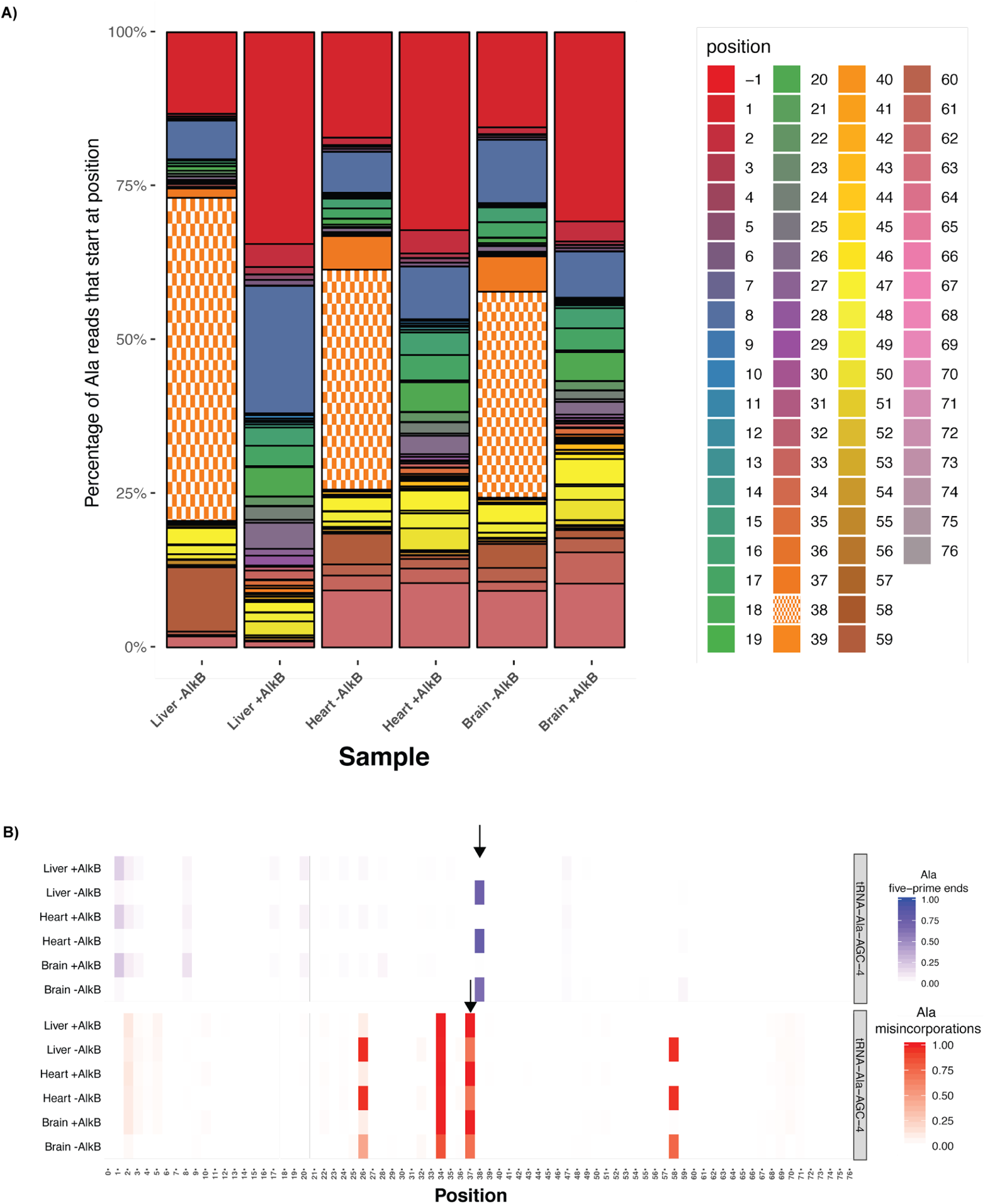
Reverse transcription (RT) terminations occur adjacent to known modification sites. **A** Percentage of sequencing reads for tRNA^Ala^ with 5’ end starting at individual positions across the length of tRNA. Position 38 (cyan) denotes the position with large propensity for RT terminations. **B** Correlation between 5’ ends of sequencing reads and detected misincorporations for tRNA-Ala-AGC-4. The top heatmap represents the percentage of reads with 5’ ends starting at positions across tRNA-Ala-AGC-4. The bottom heatmap represents the percentage of misincorporations at each position across tRNA-Ala-AGC-4. Arrows highlight position 38 for the 5’ end and position 37 for the detected misincorporation.

### Comparisons of tRAX with other tRNA-seq data analysis methods

With the discovery of tDRs in multiple organisms through small RNA sequencing, a number of data analysis methods have been developed for assessing specifically the expression of tDRs. The major ones include MINTMap [41], SPORTS [42], and tDRmapper [43]. Although these tools align sequencing reads to reference sequences and estimate abundance for tDRs similar to tRAX, they do not aim at analyzing mature or pre-tRNAs, and differ in many feature offerings and data processing methodologies (Table 1). Recently, a new method, mim-tRNAseq, was developed to study only mature tRNA abundance and modification [81]. To evaluate the performance of tRAX, we compared it with these tools by analyzing the original data sets for the ARM-seq [36] and DM-tRNA-seq [37] protocols.

**Table 1.**
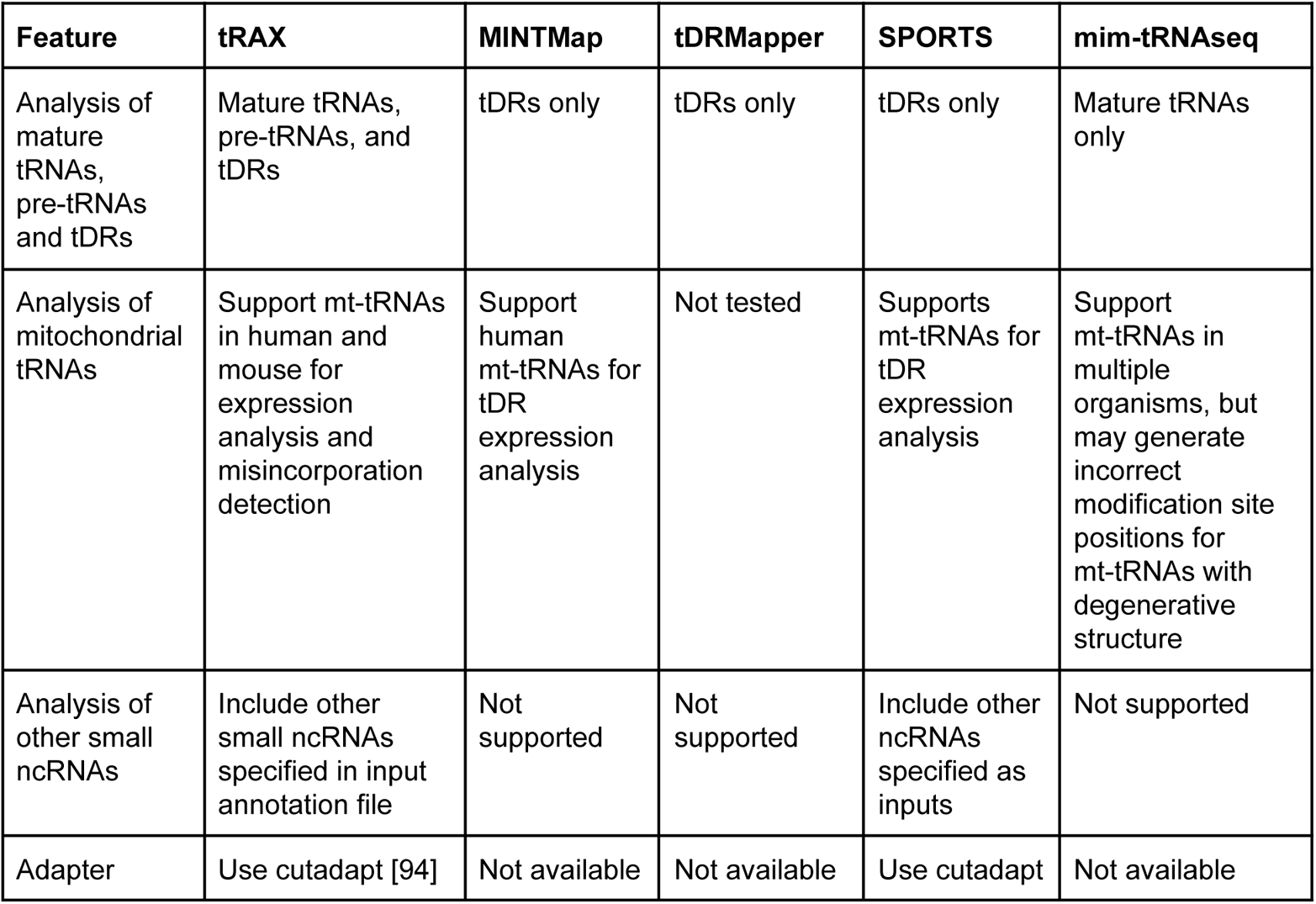
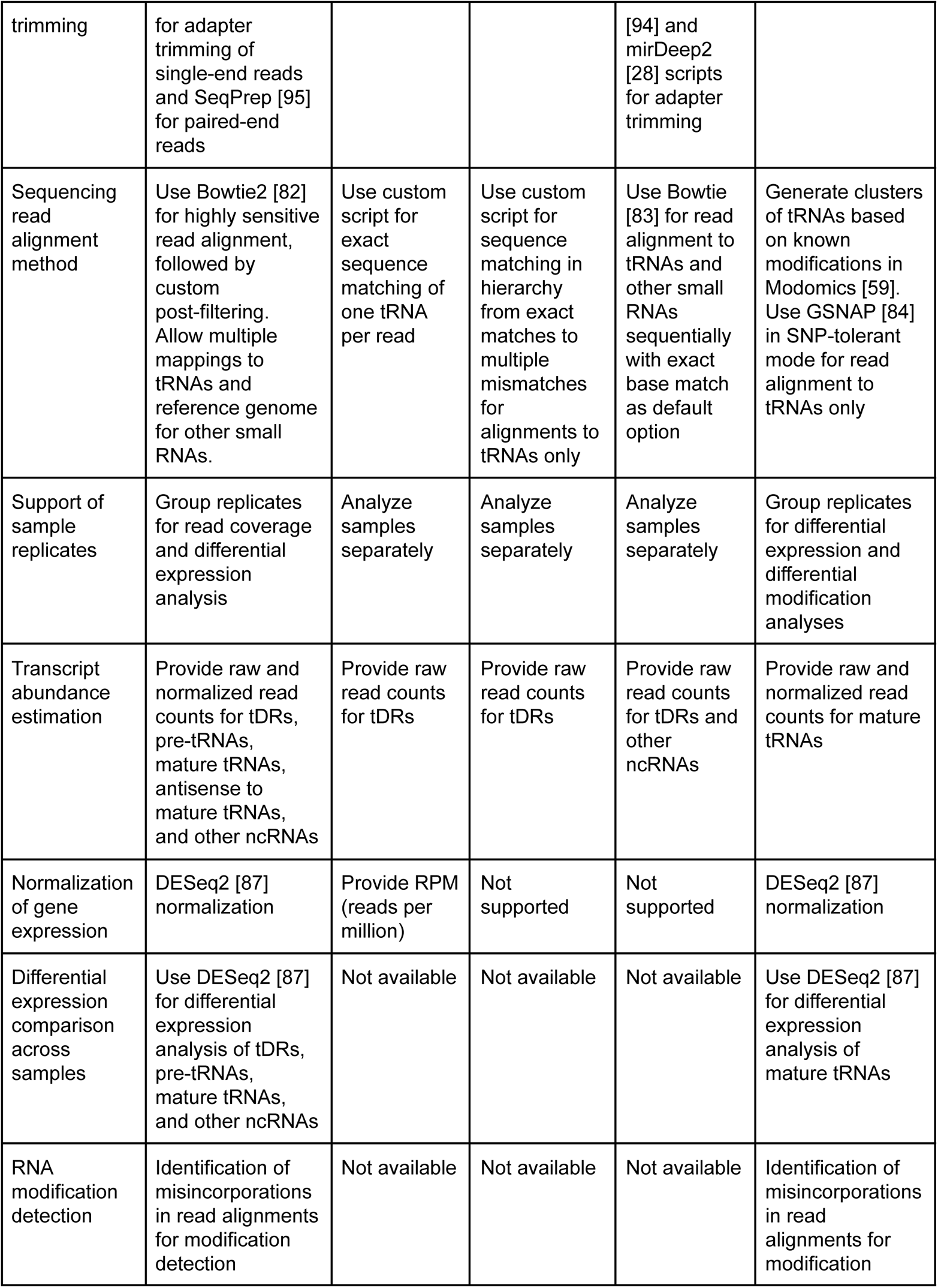
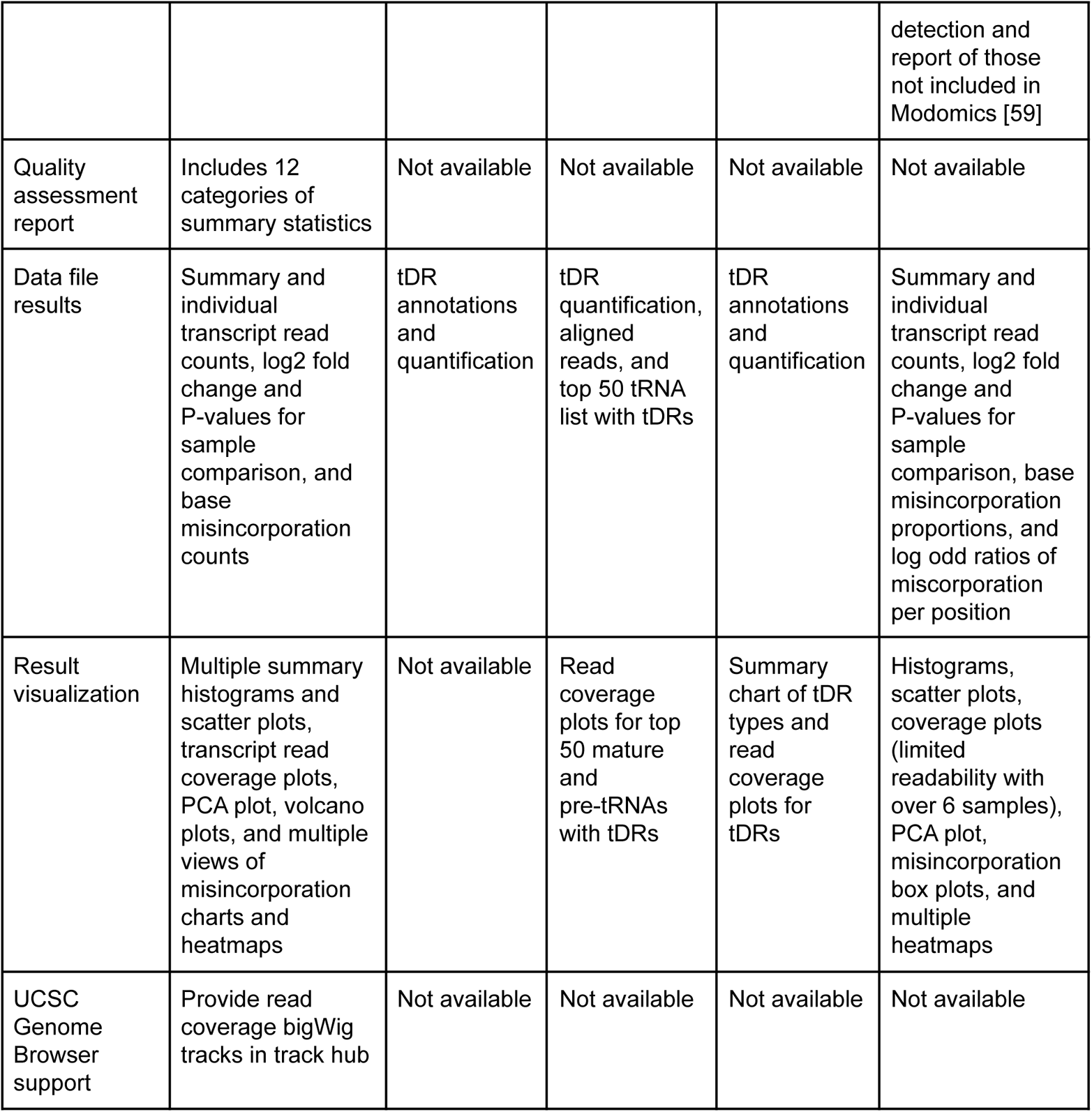
Feature comparison between tRAX and other tRNA-seq analysis pipelines

The first distinction among the five analysis packages is the availability of the data pre-processing step. Raw sequencing reads generated with small RNA-seq library preparation kits always include adapter sequences that must be removed before aligning to reference sequences. While both tRAX and SPORTS include an adapter trimming step as part of the software package, researchers are expected to pre-process the reads separately before using the other tools. All these software then map the trimmed reads to the reference sequences but with very different methods. tRAX uses Bowtie2 [82] with relaxed parameters to align reads to both tRNA transcripts and reference genome, followed by a custom post-filtering step that allows alignments at multiple loci, accommodations to misincorporations due to modifications, and also enables the determination of mappability uniqueness. (Additional File 1: Figure S19) (see Methods). Although SPORTS uses Bowtie [83], the predecessor of Bowtie2 [82], it aligns reads to tRNAs and other small RNAs sequentially, and does not allow base mismatch as a default option. On the other hand, MINTMap employs an exact-sequence-matching approach to search for partial tRNAs while tDRmapper follows a hierarchy of stages to select alignments from having exact match to multiple mismatches/deletions. Unlike tRAX, they do not support paired-end sequencing reads that are not commonly used for miRNA or tDR sequencing, but can be advantageous for sequencing pre-tRNAs and Type II tRNAs (tRNA^Leu^ and tRNA^Ser^ in all organisms; tRNA^Tyr^ also in bacteria), which are usually close to or over 100 nucleotides long. mim-tRNAseq, the newest available tool, uses GSNAP [84] with SNP-tolerant alignments based on known tRNA modifications in MODOMICS for detecting misincorporations and supports both single-end and paired-end sequencing reads. Due to the limits put into the read alignment rules, SPORTS, MINTMap, and tDRmapper reported much lower read counts than tRAX in our ARM-seq data comparison (Additional file 1: Figure S20). For example, tRAX found over 2000 reads aligned to tDRs derived from tRNA-Glu-TTC-2. Yet, MINTMap and SPORTS only reported 226 and 272 reads respectively while tDRmapper did not find any reads (Additional file 1: Figure S20, see labeled point). In spite of having high specificity to align sequencing reads to tRNAs, the sensitivity of these three packages is significantly lower than tRAX, resulting in possible missing read counts and inadequate abundance estimation of the transcripts. Although the tools do not intend to support mature tRNA analysis, we added up the read counts of detected tDRs derived from the same tRNAs to perform a comparison using DM-tRNA-seq data. Compared to the ARM-seq data, these three methods identified even fewer reads than tRAX (Additional file 1: Figure S21). Since mature tRNAs are highly modified, mismatch allowance in read alignments is an important criterion for optimally including the appropriate reads for abundance estimation. This can be observed in the results of tRAX and mim-tRNAseq that have similar read counts for most tRNAs (Additional file 1: Figure S21).

Use of sample replicates has been found to increase statistical power and accuracy when performing gene expression comparison [85, 86]. However, SPORTS, MINTMap, and tDRmapper process samples one at a time without the capability of utilizing replicates for statistical analyses. tDRmapper analysis results only include raw read counts with which researchers need to apply their own normalization methods for sample comparison. Nevertheless, it generates read coverage in text files for researchers to create charts and provides graphical plots of the read coverage for the top 50 mature and pre-tRNAs with tDRs. Although RPM (reads per million) values are computed by SPORTS as normalized transcript abundance, they are only used in the coverage plots and read length distribution charts for visualization but not included as part of the default output results. MINTMap, on the other hand, provides multiple RPM (reads per million) values for each tDR for comparison across samples. In contrast, both tRAX and mim-tRNAseq take on a more comprehensive approach to enable differential expression analysis between samples with the use of DESeq2 [87]. Similar to tRAX, mim-tRNAseq normalizes read counts with DESeq2 [87] and produces read coverage plots for each tRNA transcript; however, the fixed chart width results in reduction of readability if the study exceeds six samples. Unlike tRAX which performs comparison between all pairs of samples specified by researchers, mim-tRNAseq focuses on comparison between a specified control and other conditions. In addition, both tools generate various analyses and different collections of charts such as principal component analysis (PCA), heatmaps, and volcano plots of log_2_ fold changes against adjusted P-values for each pair of sample comparisons to assess differential expression significance. Because DESeq2 was not designed to handle experiments without sample replicates, tRAX has an option to use its predecessor DESeq instead while mim-tRNAseq skips the generation of comparison charts when no replicate is specified.

Among the five analysis tools, only tRAX and mim-tRNAseq detect possible tRNA modifications through misincorporations found in read alignments. Both tools compute the proportion of reads for each nucleotide as mismatches at each Sprinzl position of tRNA transcripts, with heatmaps to illustrate the misincorporation positions, in addition to other plots that display the misincorporation proportion and nucleotide distribution in different perspectives. mim-tRNAseq further provides a list of newly predicted modification site candidates that are not annotated in MODOMICS [59]. Yet, for the DM-tRNA-seq data set, we found that the predicted results are more accurate when combining with the provided log-odd ratios of misincorporation proportions between the AlkB-treated and untreated samples. As such, modifications such as m^1^A_58_ that have been mostly removed in the treated samples are still included in the predicted modification list. mim-tRNAseq also includes the analysis of mitochondrial tRNAs (mt-tRNAs) in addition to cytosolic tRNAs by default. However, we found that misalignments tend to result from mt-tRNAs with degenerative secondary structure [88]. On the other hand, tRAX supports mt-tRNAs by providing downloadable pre-built reference data.

In comparison with the other four tools, tRAX produces the most abundant and multi-perspective graphical images for result visualization (Additional file 2: Table S1). As described before, the read coverage plots from tRAX include the distinction between uniquely mapped reads and reads shared among tRNA isodecoders or isotypes. The use of standard tRNA Sprinzl positioning in all the per-base coverage plots allows researchers to uniformly identify tDR location and distribution of reads along tRNA transcripts. The read coverage data tracks in track hubs that can be visualized in UCSC Genome Browser provide researchers an opportunity to explore the data in genomic context. Together with the quality assessment report and multiple summary charts of read distributions which are not available from other tools, tRAX empowers researchers to examine overall data quality and diagnose potential failures in sequencing library preparations. These features have been instrumental in aiding our own group to regularly improve future experimental design.

## Discussion

We present here a unified data analysis pipeline for tRNA and tDR sequencing that offers the most complete view of tRNA biology via RNA-sequencing. It combines transcript abundance estimation, differential expression analysis at the granularity of individual isodecoders, and facile detection of RNA modifications. Furthermore, these analyses are used to generate a diverse plethora of visualizations through global-view histograms and scatter plots, as well as individual transcript read coverage profiles, misincorporation heatmaps, and genome-context views via UCSC Genome Browser data tracks, among others. By using a custom-built reference database containing tRNA transcripts and the reference genome sequence, tRAX can successfully quantify and compare expressed tRNAs, pre-tRNAs and tDRs across samples. Moreover, the algorithm design better incorporates both post-transcriptional modification and the possibility of multi-copy tRNA genes to achieve greater sensitivity and detail per transcript relative to previous tRNA sequencing analysis methods [41–43].

The addition of a quality assessment report to tRAX is a unique feature among tRNA-seq analysis software, and a necessity given the wide range of small RNA sequencing data quality and growing number of sequencing methods. While this report is not meant to be a replacement of other data quality assurance tools such as FastQC [89], it provides evaluations specifically applicable to tRNA-seq results based on issues observed in previously analyzed data sets. For example, if a particular sample or replicate does not meet the recommended criteria included in the report, researchers are alerted immediately and can consider repeating the experiment, altering sample library preparation, or making appropriate adjustments in subsequent analyses (e.g., excluding the failed replicate).

The level of detailed small RNA categorization provided by tRAX allows clearer identification of individual transcripts which show novel patterns. For example, tRAX revealed a previously undetected subset of tDRs facilitated by treatment with T4 polynucleotide kinase (Fig. 3; Additional File 1: Figure S17), which may provide additional regulatory clues to their biogenesis. Additionally, the display of all tDRs aligned to their source tRNA allows them to be studied both in the context of all that is known about the mature tRNA *and* all other derived tDRs -- this facilitates identification of common tRNA processing events and/or associated sequence motifs. Alignments use the Sprinzl [90] canonical tRNA positions (of high reference value to tRNA biologists), and include the mature tRNA secondary structure as well as annotated tRNA modifications from Modomics [59]. A unique, defining element of tRAX result visualization is the classification of reads based on uniqueness of mappability. The color-coding of sequencing read profiles into categories based on alignment specificity (transcript-, isodecoder-, or isotype-specific) allows researchers to instantly assess any issues associated with tRNA multiple mapping which is of particular importance for complex metazoan tRNA sets.

Another key innovation of tRAX is the visualization of reverse transcription (RT) misincorporation, termination, and base skipping. Early termination of primer extension has been used to identify the presence of RNA modifications for many years, but only recently has this technique been applied to high-throughput tRNA-seq data to predict specific types of modifications across the tRNA transcriptome. Visualizations from tRAX make identification of recurrent nucleotide-resolution misincorporation events for every transcript unambiguous, and plainly highlights changes between samples (e.g., tRNA-processing mutants or response to varying growth conditions). tRAX allows mapping of sequencing reads with one or more misincorporation, thus making it possible to detect multiple modifications in a single read. Future work is planned to assess and generate RT-derived sequencing “signatures” which can be applied to sequencing experiments for automated prediction of specific tRNA modifications. Currently, these modifications and per-sample changes can be manually identified with relative ease from scatter plots, heatmaps, and histograms.

When designing the program, it was a priority to make tRAX results compatible with other complementary tools. tRAX provides read alignments as standard BAM files that can be visualized in the Integrative Genomics Viewer [91] or similar tools. Most of the analysis data include different levels of summary statistics, raw and normalized read counts, log_2_ fold changes, adjusted P-values, and detected misincorporations, all provided as tab-delimited text files that can be used with spreadsheet software such as Excel or other statistical tools for further analyses. The UCSC Genome Browser tracks produced can be easily loaded as custom tracks or incorporated into a track hub. This allows the user to display read coverage data of each sample, and offer another method for studying the expression of tRNA genes in their full genomic context. These features expedite interpretation of results and the development of additional downstream analyses.

In consideration of tRAX relative to previously developed tRNA-seq analysis options, we note additional technical advantages. First, tRAX has been packaged for use with Docker (docker.com), which can eliminate the hurdles of locating and installing diverse software dependencies. tRAX is also the only tRNA-seq analysis program that includes an option to use modern multi-core computing processors for many-fold accelerated execution. tRAX is the most comprehensive of available tools, offering a complete start-to-end gene expression analysis workflow from pre-processing of raw sequencing data, mapping, and quality assessment, to differential expression analysis and visualization by more than fifty perspectives -- some of those steps, for example differential expression analyses and utilizing sample replicates, are surprisingly omitted in all previously developed tools. Furthermore, the read alignment approach that takes into account specific characteristics of tRNAs, such as identical gene copies or highly conserved isodecoders and base misincorporations due to RNA modifications, enables tRAX to deliver more accurate quantification of the transcripts in the samples. tRAX is also the only tool that was specifically designed to support expression analysis of both mature tRNAs and tDRs, allowing researchers to evaluate correlations between these two transcript types through consistent data and resources.

## Conclusions

tRAX is a comprehensive data analysis package that enables the study of tRNAs and tDRs through deep sequencing. It was developed by a tRNA biology research group with over twenty years of experience in computational and experimental analyses of tRNAs, and as such, takes into account the many complexities of tRNA transcript measurement. The authors are dedicated to long-term, in-depth support and regular improvements to tRAX as tRNA-seq methods and capabilities evolve. Because the expanding roles for tRNAs and their products have yet to be studied in broad physiological and phylogenetic contexts, the features provided by tRAX can be instrumental in delivering detailed, robust analyses in a ready-to-explore format that should spur many new advancements in tRNA biology.

## Methods

### tRAX overall design and requirements

tRAX is a multi-step pipeline for analyzing tRNA and tRNA-derived small RNA (tDR) expression profiles using high throughput sequencing. It was originally developed to support the study of tDR expression using ARM-seq protocol [92]. With the option of omitting the fragment type determination, the tool now can also work with mature tRNA sequencing protocols such as DM-tRNA-seq [37]. The software was developed using Python for the framework and data processing, R and ggplot2 for graphical chart creation, and external bioinformatics tools and data sources as dependencies that the pipeline uses in different steps as described below in more detail. It runs natively as a command-line tool on a Unix/Linux environment and can also be used via a Docker container or Conda environment in a broader context. The software was designed to be used with data generated by Illumina sequencing platform, which is the most widely used sequencing technology. We recommend researchers who would like to use it with data from other sequencing platforms to work with us on any compatibility issues. Since sequencing data files are relatively large in size, the software supports multi-thread processing and we suggest that researchers use it on a multi-core server instead of personal computers.

### Reference database for data analysis

To prevent incorrect alignment of sequencing reads, tRAX requires the use of a reference database that is specially built for tRNA and tDR analysis. A tool called maketrnadb.py is included in the tRAX package with which researchers can create the database via tRNAscan-SE [93] annotations and sequences downloaded from the Genomic tRNA database [30] along with the genome sequence of the organism. The database building tool generates a Bowtie2 read mapping index [82] that consists of both the set of unique mature tRNAs and the genome sequence of the organism. The mature tRNA sequences are created with 3′ CCA tails, removal of introns, and addition of the histidine post-transcriptional 5′ G base. As an additional step, 20 “N” bases flanking to each side of the sequences are added to allow for extra bases off the end of the tRNAs such as potential CCACCA ends. Moreover, alignments of both mature tRNA sequences and genomic tRNA sequences are generated using cmalign of Infernal v1.1 [88] with options “--nonbanded --notrunc -g” and tRNA covariance models from tRNAscan-SE [93] for computation of per-base read coverage and misincorporation frequencies. To increase the accuracy of the alignments, a sequence source option (--orgmode) can be specified when generating the reference database for using covariance models trained with tRNAs in the corresponding clade. The current version supports cytosolic tRNAs in eukaryotes with the expansion to include archaeal and bacterial tRNAs in development.

### Adapter trimming and paired-end read merging

The first step of analyzing small RNA sequencing data is the removal of adapters used in sequencing library preparation from raw sequencing reads. tRAX includes a separate adapter trimming tool called trimadapters.py that researchers can adapt or select to use other preferred methods for the same task. This adapter trimming tool uses cutadapt [94] to remove the adapter sequences from single-end reads that are commonly employed for small RNA sequencing. If the length of the sequencing reads is shorter than the minimum required read length, which is 15 nt as default, those reads will not be retained for the next step to avoid large amounts of ambiguous read alignments. For paired-end reads, the sequencing inserts which are small RNA transcripts usually “overlap” at the two read ends because the read lengths are mostly longer than the transcript lengths. Bowtie2 [82] that is used in tRAX for read alignment cannot correctly align “overlapping” read ends to the reference sequences. Therefore, tRAX uses a tool called SeqPrep [95] that can remove the adapter sequences and merge the two read ends into pseudo-single-end reads at the same time. The default adapter sequences to be trimmed are “AGATCGGAAGAGCACACGTC” and “GATCGTCGGACTGTAGAACTC” that are mostly used in small RNA-seq library preparation kits for the Illumina sequencing platform. Researchers who use custom sequencing library preparation methods can specify optionally the desired adapter sequences. The sequencing reads retained after trimming can be used for read mapping in the following step of the process and the summary statistics of adapter trimming are included in the quality assessment report described below.

### Aligning sequencing reads to reference database

Sequencing reads without adapters should be used as inputs to the main tRAX tool, processsamples.py, that includes read mapping, abundance estimation, differential expression analysis, and misincorporation estimation for RNA modification identification. Sequencing reads are aligned to the pre-built reference database using Bowtie2 [82] in very-sensitive mode that ignores quality scores and allows a maximum of 100 alignments per read (--very-sensitive --ignore-quals --np 5 -k 100). Due to the presence of modifications and editing in tRNAs, tRAX does not use a hard mismatch cutoff by default. Instead, read alignments are post-filtered to include only all best mappings, with an exception that when reads map equally well to mature tRNAs and genome sequence, those mapped to the genome are removed. Using this method, reads that contain genomic sequence flanking the tRNA loci in the genome will be considered to be part of the pre-tRNAs, while reads that contain no flanking sequence will be considered to be derived from mature tRNAs. If the “primary” alignment defined by Bowtie2 [82] is not included in the selected alignment set, another alignment will be chosen arbitrarily. Reads that do not align to tRNA transcripts but to genomic sequences and are less than 16 bases in length are discarded to avoid high rate of off-target alignments. This custom read mapping method allows mismatches in alignments that may represent misincorporations due to RNA modifications while keeping most of the reads uniquely aligned to single transcripts (Additional File 1: Figure S19). Sorted and indexed BAM files are generated that will be used for abundance estimation or visualization in alignment viewers such as Integrative Genomics Viewer [91].

### Abundance estimation of tRNAs, tRNA fragments, and other small RNAs

tRAX computes read counts for both tRNAs and other small RNA genes that are supplied by the user. The non-tRNA gene annotations have to be specified in Ensembl GTF file format with the classification of gene types. Sequencing reads that align to more than 10 nt of the mature tRNA sequences or other small RNA genes are included in the raw read counts of the corresponding genes. Reads that align to the antisense of mature tRNA sequences are included separately. Additionally, when analyzing tDRs, tRAX separates the sequencing reads that map to mature tRNAs into four fragment types: whole tRNAs, 5′ fragments, 3′ fragments, and “other” fragments (Additional file 1: Figure S10 – S13). The fragment types are determined by the distance between the read alignments and the ends of the tRNA. Reads where both the 5′ end and 3′ end lie within 5 nt of their respective ends on the mature tRNAs are categorized as “whole tRNAs”. Reads that only overlap or align closely to the 5′ end of the tRNAs are considered as “5′ fragments”. Similarly, reads that only overlap or align closely to the 3′ end of the tRNAs are “3′ fragments”. Fragments that do not meet any of these criteria are “Other fragments”. The 5′ and 3′ fragment types loosely correlate to the known TRF-5 and TRF-3 fragment types [96] but the requirements are relaxed to account for the diversity of fragment types seen in ARM-seq and other sequencing experiments. This fragment classification scheme is intended to separate TRF-5 and TRF-3 tDRs that have been previously described [97], but also allow for new types of tDRs to be counted. Since pre-tRNA transcripts may be included in the sequencing libraries, tRAX expects those reads to contain flanking sequences of the mature tRNAs. Specifically, they must include at least 3 bases upstream of the gene or at least 5 bases downstream to be considered pre-tRNA fragments. Pre-tRNA reads are categorized into three types analogous to the fragment type: “whole pre-tRNAs” that start upstream and end downstream of the mature tRNAs, “partial pre-tRNAs” that overlap only part of the mature tRNAs, and “tRNA trailers” that start downstream of the mature tRNAs, corresponding to tRF-1 fragments [27]. The raw read counts are provided as a tab-delimited file that is used for normalization and differential expression analysis.

To provide summary statistics on read alignment distributions, tRAX computes the total read counts aligned to each type of genes in the provided samples. Only genes with at least 10 read alignments are included. The gene type categorization is based on the feature included in the user-specified gene annotation GTF file. To prevent double-counting, only the primary read alignment is used and each read can only be assigned to one gene, unlike the read counting step where multiple mapping is allowed. tRAX outputs text files and histograms to display the distribution of read counts for each gene type and tRNA isotype in each sample (Fig. 1B-C, and Additional file 1: Figure S2, S3, S8). In addition, the distribution of read length is calculated and presented in corresponding text file and chart with comparison between tRNAs and other gene types (Additional file 1: Figure S4 and S5).

### Normalization of read counts and differential expression analysis

Raw read counts of tRNAs, tDRs, and other small RNAs are combined as input to DESeq2 [87] for computing the normalization size factors of each sample which, in turn, are used for calculating the normalized read counts to allow comparison across samples. tRAX takes in metadata of experimental design that includes groups of replicates for the same tissue type, treatment, or condition, and sample pairs for expression comparison. In addition, DESeq differential expression pipeline from DESeq2 [87] is used with parameter “betaPrior=TRUE” for generating log2 fold changes and adjusted P-values for each specified sample pair. Normalized read counts, log2 fold changes, and adjusted P-values are provided in tab-delimited files. Researchers can also visualize the expression comparison between samples through volcano plots of log2 fold changes against adjusted P-values (Additional file 1: Figure S6 and S7). Moreover, to help review the correlation between replicates, principal component analyses of first versus second dimensions using prcomp in R are performed for all gene types in the study and only tRNA/tDRs.

### Per-base read coverage for tRNAs and tRNA fragments

Using the read alignments, tRAX determines the start and end positions of sets of reads aligned to tRNAs and tRNA fragments. The region in between the start and end positions is considered to have read coverage. Per-base coverage is computed with the Sprinzl positions of tRNAs, the canonical positioning of all tRNA transcripts [90], which is used in the tRNA alignments pre-built in the reference database (see above). The coverage values in each sample are then normalized with the corresponding size factor generated using DESeq2 [87] and provided in tab-delimited files. The values are further averaged across replicates of the same tissue/treatment and displayed as combined and individual graphic coverage plots for each tRNA isodecoder (Fig. 2D–F, Additional file 1: Figure S9, S19). To assess the uniqueness of reads aligning to tRNAs, tRAX separates the read coverage calculation into reads uniquely mapped to gene loci, reads aligned to multiple tRNA isodecoders, reads aligned to multiple tRNAs with different anticodons of the same isotype, and reads aligned to multiple tRNA isotypes. Different colors are used in the coverage plots to represent the different read mapping categories (Additional file 1: Figure S21).

To allow researchers visualizing the expression of tRNAs and tRNA fragments in genomic context, tRAX generates a UCSC Genome Browser track hub [51, 98] that contains bigWig tracks for the read coverage of each sample. Specific BAM files are created based on the original ones from the step of aligning reads (see above) in which reads are now assigned to the genomic tRNA loci of the corresponding isodecoders with adjustment of genomic coordinates over introns if existed. genomeCoverageBed from Bedtools [99] is used to convert these BAM files into read coverages in bedGraph file format that are further converted into the bigWig files using bedGraphToBigWig in UCSC Genome Browser Utilities [100].

### Misincorporation and deletion detection for tRNAs

As RNA modifications in tRNAs can be detected by misincorporations in sequencing reads using DM-tRNA-seq with TGIRT reverse transcriptase [78], tRAX computes the mismatch and deletion frequencies observed in the read alignments. Similar to the per-base coverage computation, read alignments are piled up according to Sprinzl positions of tRNAs where tRAX counts the occurrence of each nucleotide and deletion with comparison to the reference sequences. The raw counts of mismatches and deletions are normalized using the DESeq2 [87] size factors of each sample. A series of graphical images including dot plots, bar charts, and per-base histograms by Sprinzl positions, tRNA isodecoders, anticodons, and isotypes are produced to represent the mismatch and deletion frequencies (Fig. 5 and 6B). For the per-base histograms, a pseudocount of 20 reads is used to reduce the visibility of low-coverage reads. In addition, mismatch frequencies are presented in heatmaps for each tRNA isodecoder using a pseudocount of 20 for mismatches.

### Generation of quality assessment report

tRAX generates a quality assessment report in HTML format to provide researchers with an estimate on the “quality of data” and possible diagnostic information when issues occur. Eleven quality metrics are included to determine the data quality at different processing stages (Table 2).

**Table 2.** Quality metrics in quality assessment report for determining the sequencing data quality

| Category | Fragment mode | Non-fragment (full length tRNA) mode |
| --- | --- | --- |
| Sequencing read filtering/merging rate (percentage of sequencing reads retained for read mapping) | pass: $\geq 60\%$; warning: $< 60\%$ and $\geq 20\%$; fail: $< 20\%$ | pass: $\geq 60\%$; warning: $< 60\%$ and $\geq 20\%$; fail: $< 20\%$ |
| Number of mappable reads | pass: $\geq 200,000$;<br>warning: $< 200,000$ | pass: $\geq 200,000$;<br>warning: $< 200,000$ |
| Read mapping rate | pass: $\geq 65\%$;<br>warning: $< 65\%$ | pass: $\geq 65\%$;<br>warning: $< 65\%$ |
| Distribution of reads mapped to tRNAs | pass: $\geq 50\%$;<br>warning: $< 50\%$ | pass: $\geq 50\%$;<br>warning: $< 50\%$ |
| Distribution of reads mapped to rRNAs | pass: $\leq 35\%$;<br>warning: $> 35\%$ | pass: $\leq 35\%$;<br>warning: $> 35\%$ |
| Distribution of reads mapped to unannotated regions | pass: $\leq 35\%$;<br>warning: $> 35\%$ | pass: $\leq 35\%$;<br>warning: $> 35\%$ |
| Optimal average read length | pass: $\leq 40$ nt;<br>warning: $> 40$ nt | pass: $\leq 75$ nt;<br>warning: $> 75$ nt |
| Read length range between 40 and 75 nt | Not applicable | pass: $\geq 70\%$;<br>warning: $< 70\%$ |
| Read length range between 15 and 50 nt | pass: $\geq 70\%$;<br>warning: $< 70\%$ | Not applicable |
| Percentage of tRNAs having at least 20 mapped reads | pass: $\geq 50\%$;<br>warning: $< 50\%$ | pass: $\geq 50\%$;<br>warning: $< 50\%$ |
| DESeq2 [87] normalization size factor differences across samples | pass: $< 3X$;<br>warning: $> 3X$ | pass: $< 3X$;<br>warning: $> 3X$ |

### RNA isolation from mouse tissues

Isolation of total RNA from whole brain, heart and liver tissue of three wildtype male C57BL/6JJcl mice at sexually mature developmental stage was performed using Direct-Zol RNA MiniPrep Kit (Zymo Research) with TRI Reagent (Molecular Research Center, Inc.), typically yielding 400–450 μg of total RNA. Manufacturer’s recommended volume of TRI Reagent was added to each tissue sample, along with ~400 uL of zirconium silicate beads (1.0 mm), and samples were homogenized using a tissue homogenization centrifuge. 50 μg of total RNA from each tissue was processed using the MirVana miRNA Isolation Kit (Life Technologies), according to the manufacturer’s instructions, to select for RNA <200 nt. RNA was concentrated using a RNA Clean and Concentrate-25 (Zymo Research), after an on-column DNase I treatment to remove residual DNA (New England Biolabs). Small RNA samples were divided and treated as minus AlkB control treatment and plus-AlkB experimental treatment. AlkB treatment was performed as previously described [36]. Finally, RNA was concentrated after each enzymatic treatment using an RNA clean and concentrator-5 kit (Zymo Research). Treated RNA was used for library preparation. RNA 3’ de-phosphorylation was carried out as previously described [101]. Briefly, AlkB-treated small RNAs were subsequently treated with T4 Polynucleotide Kinase (T4PNK; New England Biolabs) in a modified 5x reaction buffer (350 mM Tris-HCl, pH 6.5, 50 mM MgCl2, 5mM dithiothreitol) [101] under low pH conditions in the absence of ATP for 30 mins. The reaction was stopped by acid phenol/chloroform extraction for RNA isolation. For 5’ phosphorylation of small RNAs, AlkB-treated, PNK-treated (for 3’ de-phosphorylation) RNA was then subjected to additional T4PNK treatment under manufacturer’s suggested conditions for 30 mins.

### ARM-seq library preparation

For ARM-seq, libraries were constructed as previously described [92] which utilizes the NEBNext Multiplex Small RNA Library Prep Set (New England Biolabs). Treated RNA (Minus- or Plus-AlkB treatment; 1μg total small RNA) was used as input into the library preparation. All PNK-treated libraries used 500 ng of total small RNA. PCR-amplified libraries were purified using DNA Clean and Concentrate-5 (Zymo Research) and then size-selected (140-250 nts) on a 6% non-denaturing TBE-acrylamide gel. Libraries were eluted from gel pieces using Gel Elution Buffer (New England Biolabs) and precipitated using 0.3 M NaOAc, 80% ethanol, and 1 μL of Linear Acrylamide (supplied in NEBNext Kit) at final concentration. Samples were left in −80°C overnight to precipitate. Precipitated libraries were then pelleted, washed twice in 80% ethanol, and resuspended in pure H_2_O. Libraries were then quantified using NanoDrop and Agilent DNA High Sensitivity kit. One liver sample failed library preparation, so only two liver samples were used for downstream analyses.

### DM-tRNA-seq library preparation

For DM-tRNA-seq, libraries were constructed using the TGIRT™ Improved Modular Template-Switching RNA-seq Kit (InGex, LLC), which utilizes Illumina-compatible adapters for amplification and sequencing purposes. Libraries were amplified using PCR primers supplied with the NEBNext Small RNA library Prep Set (New England Biolabs) and purified using Agencourt AMPure XP beads (Beckman). Resulting purified sequencing libraries were quantified using NanoDrop and Agilent DNA High Sensitivity kit. Libraries were then pooled in equimolar amounts and concentrated using a DNA Clean and Concentrate-5 (Zymo Research). Pooled PCR-amplified libraries were size-selected (140-250 nts) on a 6% non-denaturing TBE-acrylamide gel to remove unwanted primer dimer products. Libraries were eluted from gel pieces using Gel Elution Buffer (New England Biolabs) and precipitated using 0.3 M NaOAc, 80% ethanol, and 1 μL of Linear Acrylamide (supplied in NEBNext Kit) at final concentration. Samples were left in −80°C freezer overnight to precipitate. Precipitated libraries were then pelleted, washed twice in 80% ethanol, and resuspended in pure H_2_O.

### RNA sequencing and analysis of mouse samples

Libraries prepared for ARM-seq and DM-tRNA-seq were sequenced using Illumina NextSeq. 75-nt pair-ended reads were produced as FASTQ files that were analyzed using tRAX. Sequence adapters were trimmed and pair-ended reads were merged using the tool trimadapters.py in tRAX. Reference database for tRAX was built with high-confidence tRNA predictions retrieved from the Genome tRNA Database [30] and the sequences of *Mus musculus* genome assembly GRCm38. Other gene annotations were obtained from Ensembl release 99. Biological replicates of each tissue type were grouped together as sample replicates for tRAX inputs and different tissue types were marked as pairs for differential expression comparison. For ARM-seq samples, default options of tRAX were used while --nofrag option was employed for DM-tRNA-seq samples to skip the tRNA fragment determination.

### Computation method for comparison of tRNA-specific analysis pipelines

ARM-seq data and DM-tRNA-seq data used for the pipeline comparison were retrieved from previous studies [36, 37]. Sequencing reads were preprocessed using cutadapt [94] to remove adapters from single-end reads or SeqPrep [95] to remove adapters and merge paired-end reads. A minimum read length cutoff of 15 bases was used for both methods. Annotated human tRNA sequences were obtained from the Genomic tRNA Database (GtRNAdb) [30]. Both GRCh37 and GRCh38 human genome assemblies were used as reference genomes for separate tRAX runs. Ensembl releases 75 and 96 gene sets were used respectively with GRCh37 and GRCh38 genomes. For comparing tRAX to other sequencing pipelines in ARM-seq data analysis, the untreated GM05372 sample was utilized, while the AlkB-treated sample was used for DM-tRNA-seq data comparison. tRAX default options were used for ARM-seq data analysis and --nofrag option was employed for DM-tRNA-seq data analysis. SPORTS 1.0 [42] was run with the -k option on the ARM-seq data. For DM-tRNA-seq data, -k option and the maximum sequence size extended to 200 bases were used. MINTmap [41] and tDRMapper [43] were executed with default parameters on both data sets. The GRCh38 genome was used for SPORTS while the GRCh37 genome was used for MINTmap and tDRMapper due to the unavailability of the prebuilt reference data for GRCh38. mim-tRNAseq which was designed for studying mature tRNAs was only employed for analyzing DM-tRNA-seq data. Execution parameters (--species Hsap --cluster --cluster-id 0.95 --snp-tolerance --cca-analysis --threads 15 --min-cov 2000 --max-mismatches 0.1 --max-multi 4 --remap --remap-mismatches 0.075) suggested for sample dataset provided in the software package were used for the analysis.

## Declarations

### Availability of data and materials

tRAX is open source and publicly available under GPL v3.0 license at http://trna.ucsc.edu/tRAX/ with the source code downloadable at https://github.com/UCSC-LoweLab/tRAX. The sequencing data and expression profiles of mouse brain, heart, and liver were submitted to NCBI GEO database with accession numbers: GSE140099, GSE140098, and GSE140096. ARM-seq and DM-tRNA-seq data used for tool feature comparison were downloaded from NCBI SRA accession numbers SRP056032 and SRP055858 respectively. The complete collection of all output files generated by tRAX for this study can be downloaded at the tRAX website listed above.

## Acknowledgements

We would like to thank Aaron Cozen and Eva Hrabeta-Robinson, previously of the Lowe Lab, and Eric Phyzicky and the Phizicky Lab (University of Rochester), for their feedback and original work on development of the ARM-seq protocol, which fostered initial development of tRAX. We also would like to thank Alex Bagi for feedback and packaging tRAX for use with Docker. Additionally, we thank Aidan Manning for feedback and work on development of additional tRAX visualization options with DM-tRNA-seq.

## Authors’ contributions

AH, PC, and TL defined the data analysis feature requirements. AH designed and developed the analysis methods. AH and JH performed the sequencing experiments and analyzed the data. AH and PC performed software tool comparison analysis. All authors wrote and edited the manuscript. The authors read and approved the final manuscript.

## Ethics approval and consent to participate

Not applicable.

## Consent for publication

Not applicable.

## Supplementary information

Additional File 1: Supplementary Figures

Additional File 2: Supplementary Table S1

Additional File 3: Example quality assessment report

Additional File 4: Supplementary Table S2 – S4

