## Supplemental Figures for "tRNA Analysis of eXpression (tRAX): A tool for integrating analysis of tRNAs, tRNA-derived small RNAs, and tRNA modifications"

### Supplementary Figures

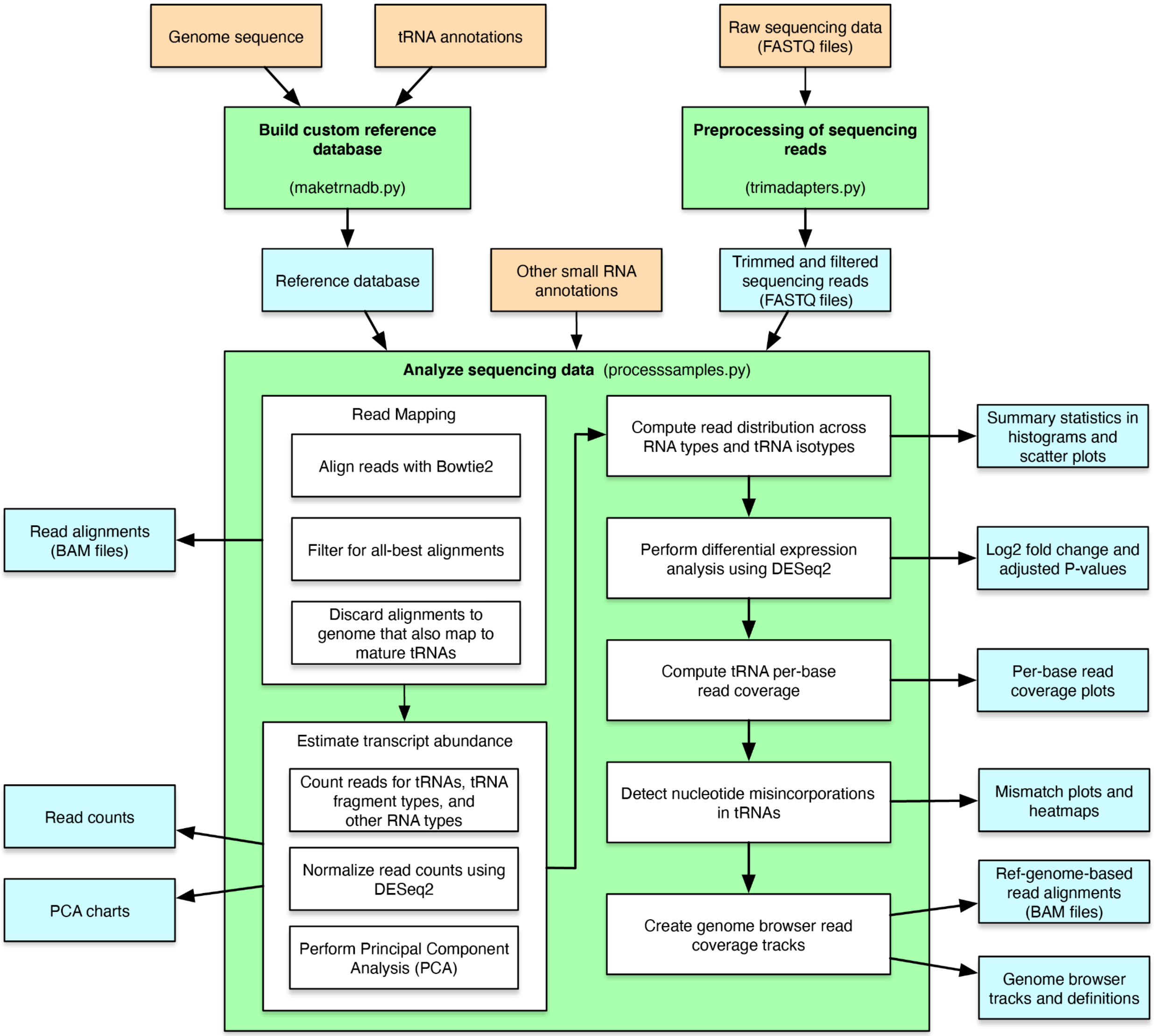


**Figure S1** Workflow of tRAX. Schematic diagram of tRAX data analysis process. It includes building of a custom reference database from user-provided genome sequence and tRNA annotations, preprocessing of sequencing data, read mapping, transcript abundance estimation, differential expression analysis, detecting misincorporations for RNA modification prediction, and creation of genome browser data tracks. Boxes in orange represent input data provided by users. Boxes in green represent the three main process modules in tRAX with the command names included in parentheses. Boxes in blue represent the output results generated by tRAX.

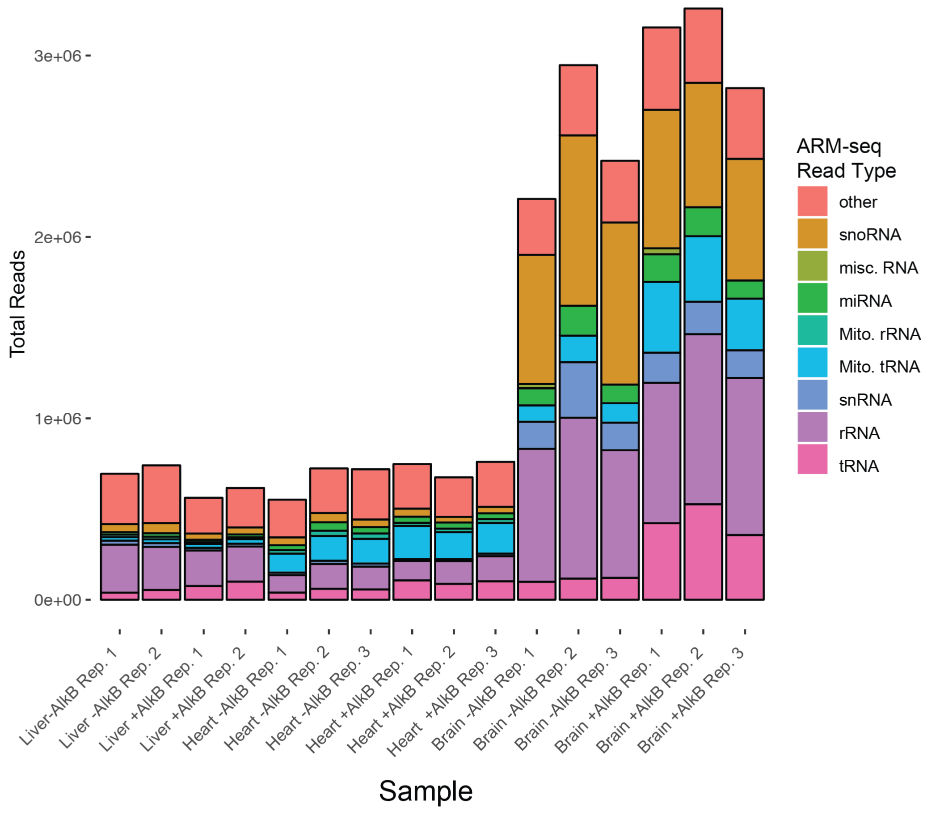


**Figure S2** Mouse tissue ARM-seq total read counts by RNA type**.** Bar graph shows the raw read counts categorized by RNA type for each mouse tissue ARM-seq library replicate. +/- AlkB represents with or without AlkB treatment.

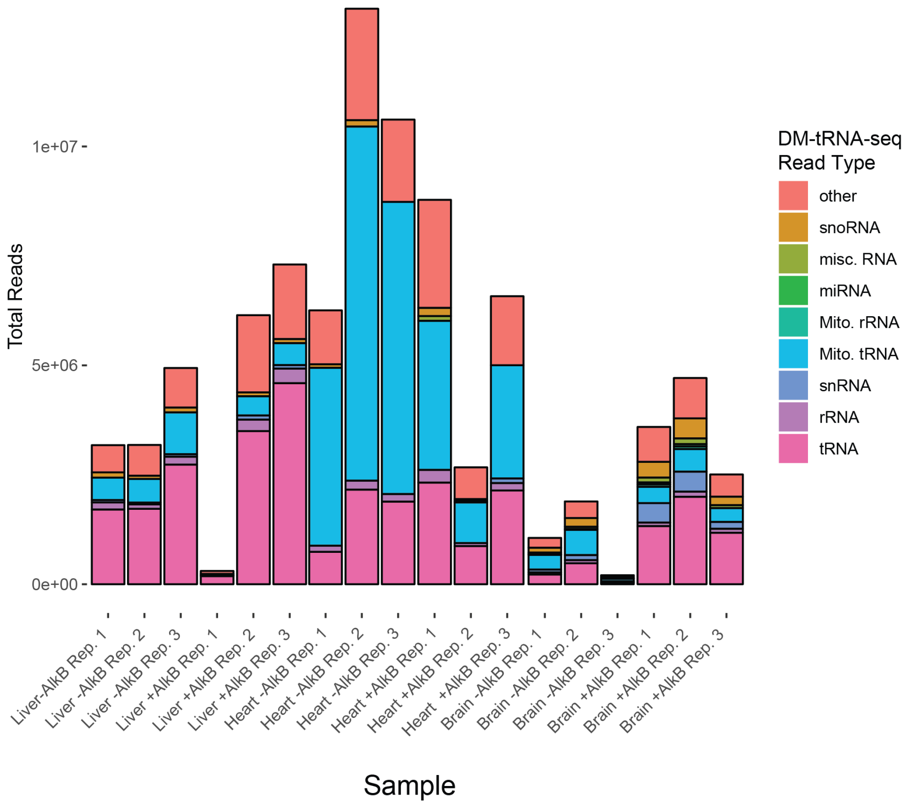


**Figure S3** Mouse tissue DM-tRNA-seq total read counts by RNA type. Bar graph shows the raw read counts categorized by RNA type for each mouse tissue DM-tRNA-seq library replicate. +/- AlkB represents with or without AlkB treatment.

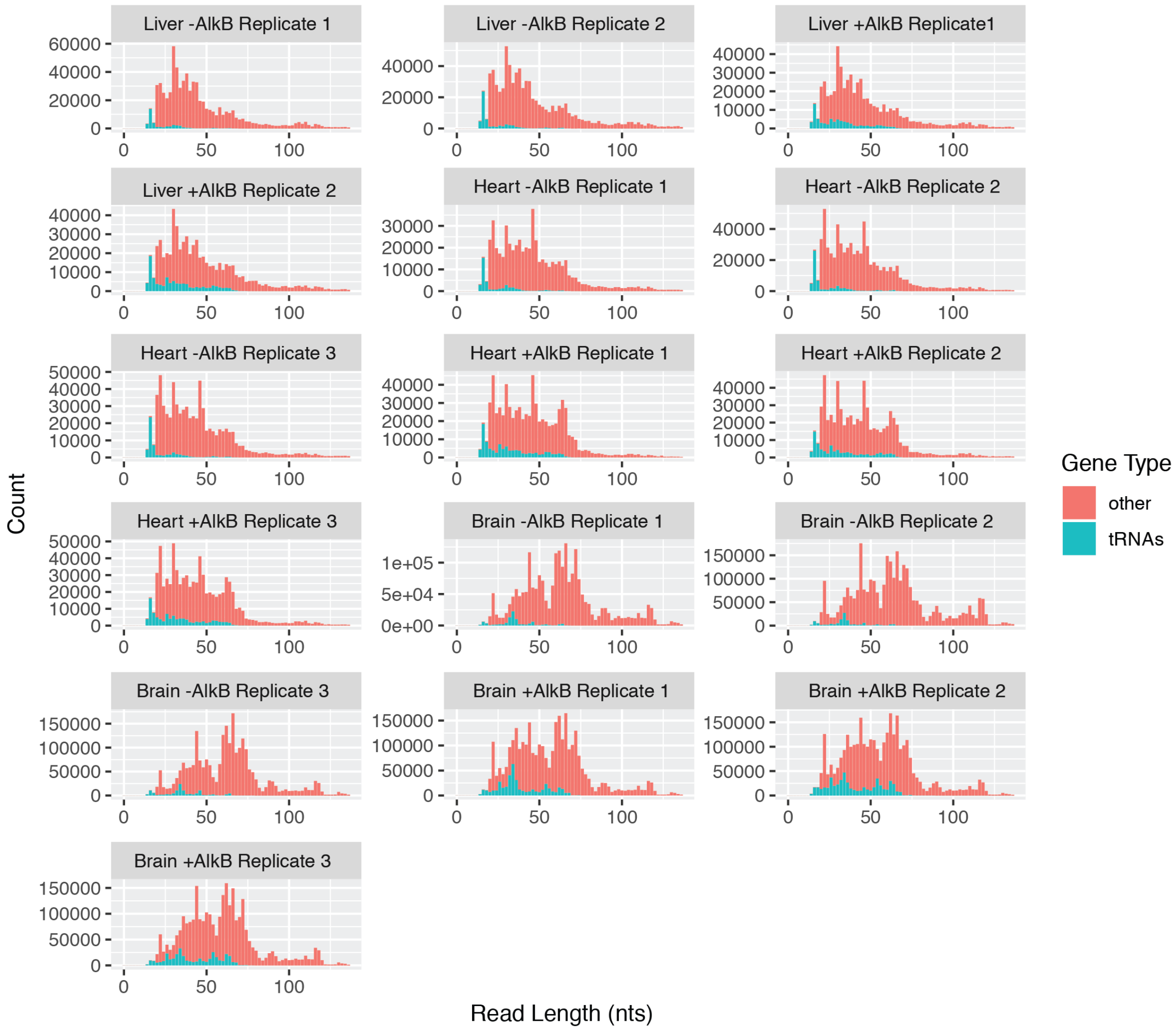


**Figure S4** Read length distribution for ARM-seq library replicates by RNA type. Bar graphs depict the read length of all sequencing reads aligned to tRNA genes ("tRNAs", *cyan*) and all other RNA types ("other", *red*). +/- AlkB represents with or without AlkB treatment. The increase in tRNA read length and abundance is visible in AlkB+ samples, and varies by tissue.

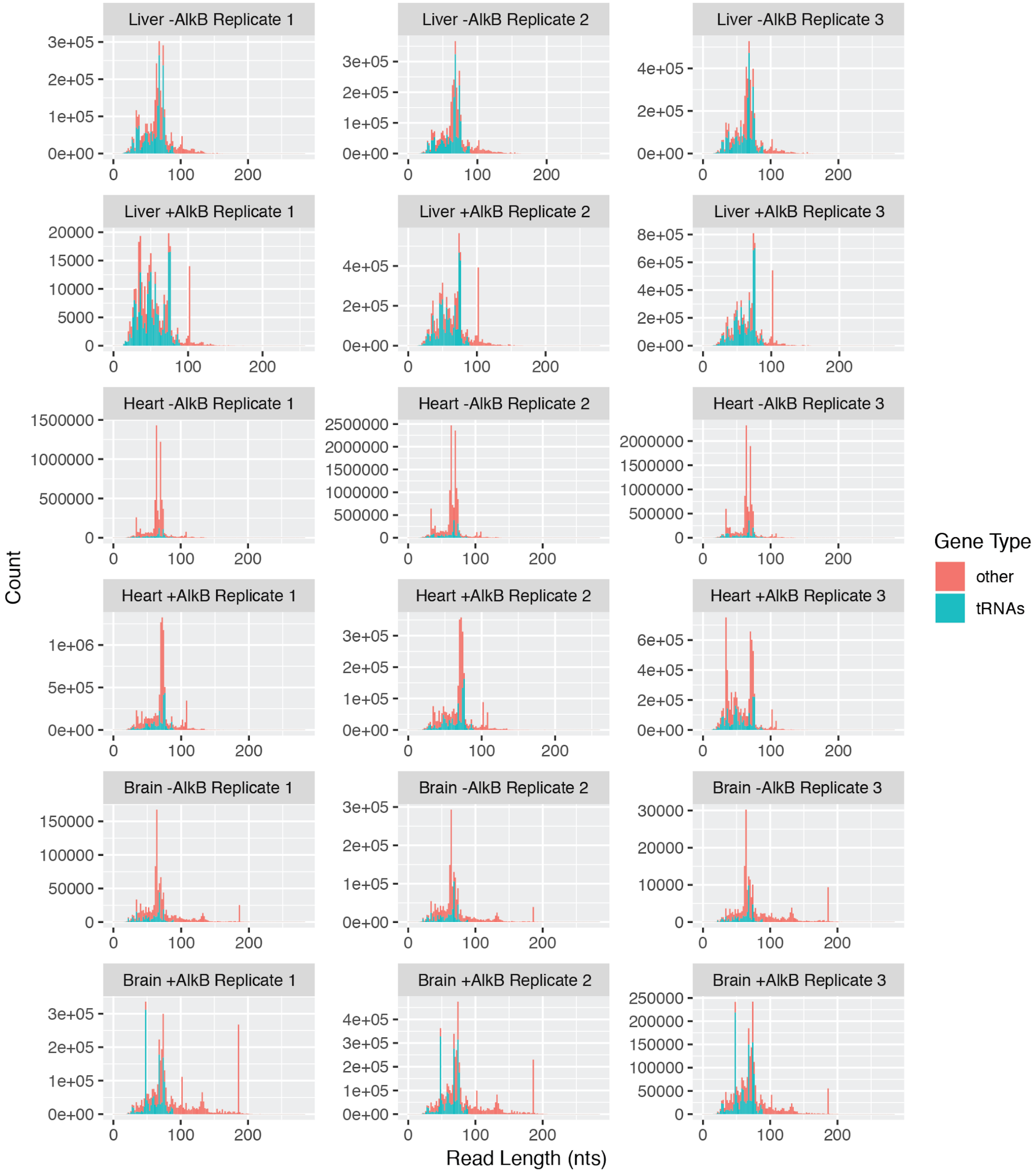


**Figure S5** Read length distribution for DM-tRNA-seq library replicates by RNA type. Bar graphs depict the read length of all sequencing reads aligned to tRNA genes ("tRNAs", *cyan*) or all other RNA types ("other", *red*). +/- AlkB represents with or without AlkB treatment. The increase in tRNAs relative to other RNA types is clearly visible in AlkB+ samples, and varies markedly by tissue.

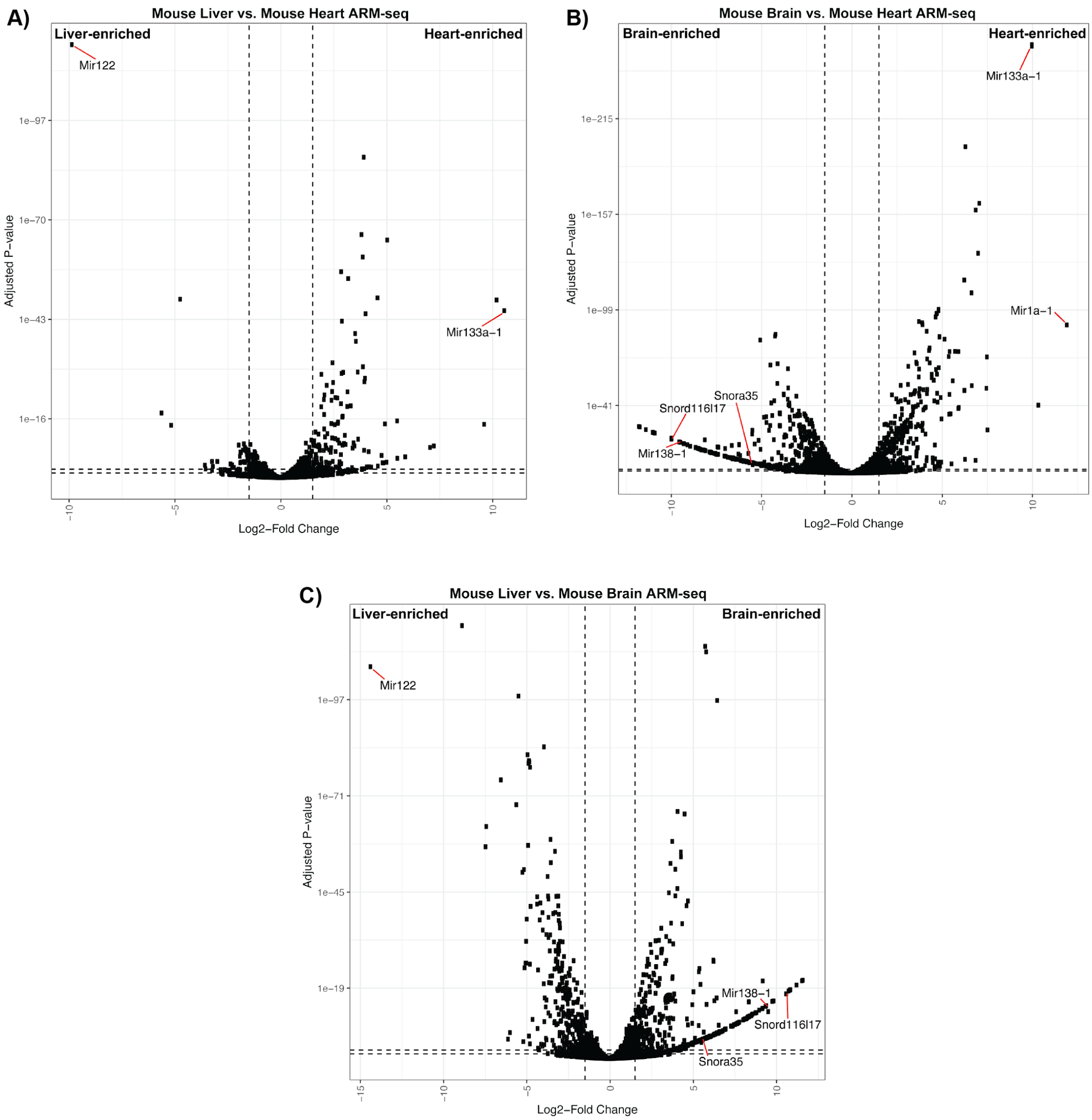


**Figure S6** Volcano plots of ARM-seq differential expression between mouse brain, liver, and heart. $Log_2$ fold changes and adjusted p-values of (A) liver vs heart, (B) brain vs heart, and (C) liver vs brain are provided. Example RNAs highlighted in Fig. 1d are called out here for direct illustration. Vertical dotted lines represent reference $log_2$ fold change values of -/+ 1.5, while horizontal dotted lines are reference adjusted p-values of 0.05 and 0.005.

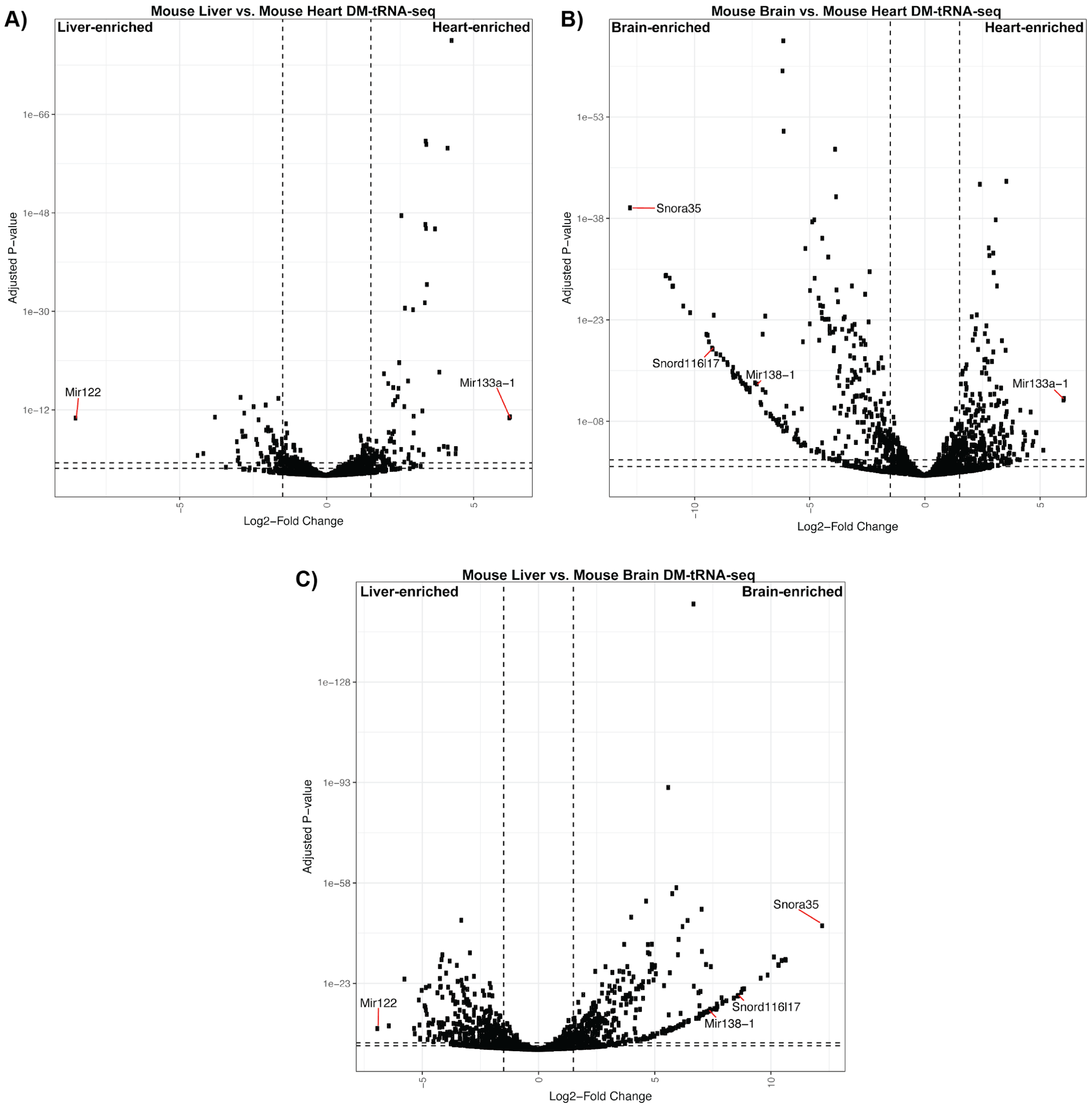


**Figure S7** Volcano plots of DM-tRNA-seq differential expression between mouse brain, liver, and heart. $Log_2$ fold changes and adjusted p-values of (A) liver vs heart, (B) brain vs heart, and (C) liver vs brain are provided. Example RNAs highlighted in Fig. 1e are called out here for direct illustration. Vertical dotted lines represent reference $log_2$ fold change values of -/+ 1.5, while horizontal dotted lines are reference adjusted p-values of 0.05 and 0.005.

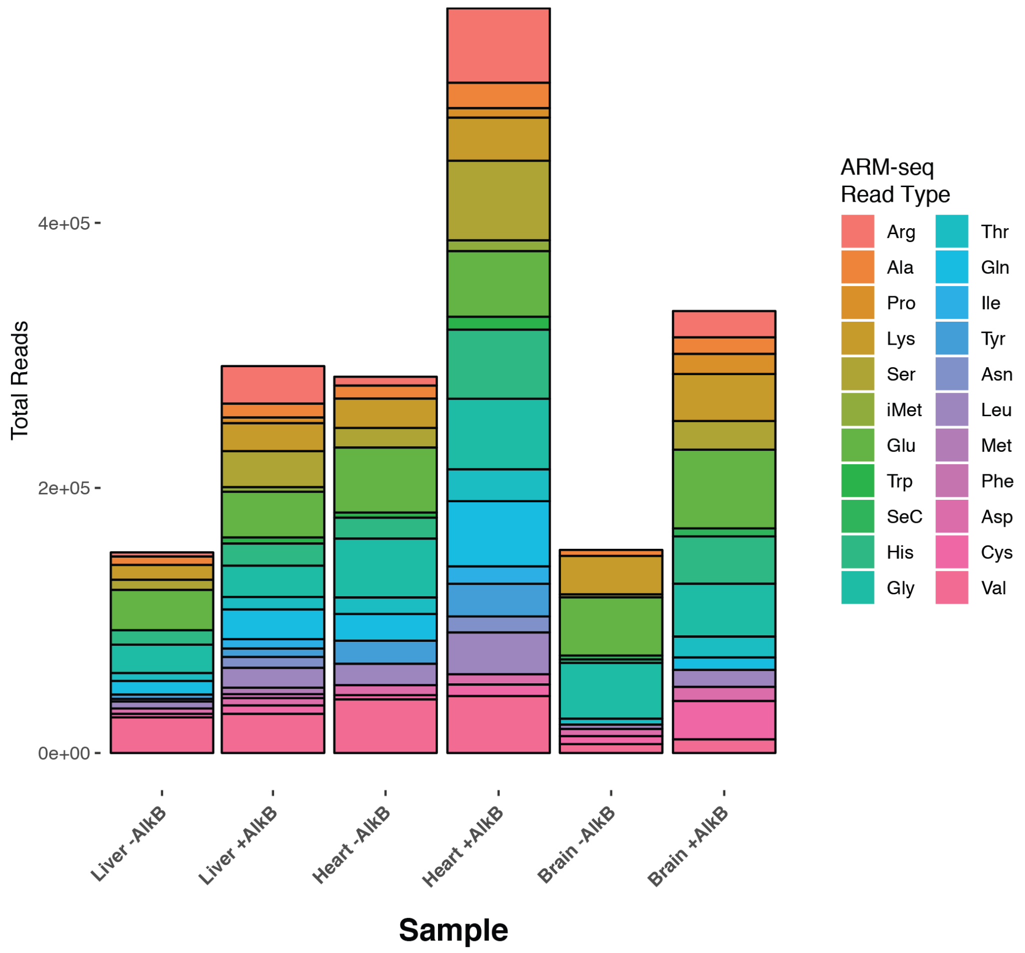


**Figure S8** Total read count by tRNA isotype for mouse tissue ARM-seq experiments. Bar graph shows the total number of reads mapped to different tRNA isotypes for all samples with (+) and without (-) AlkB treatment.

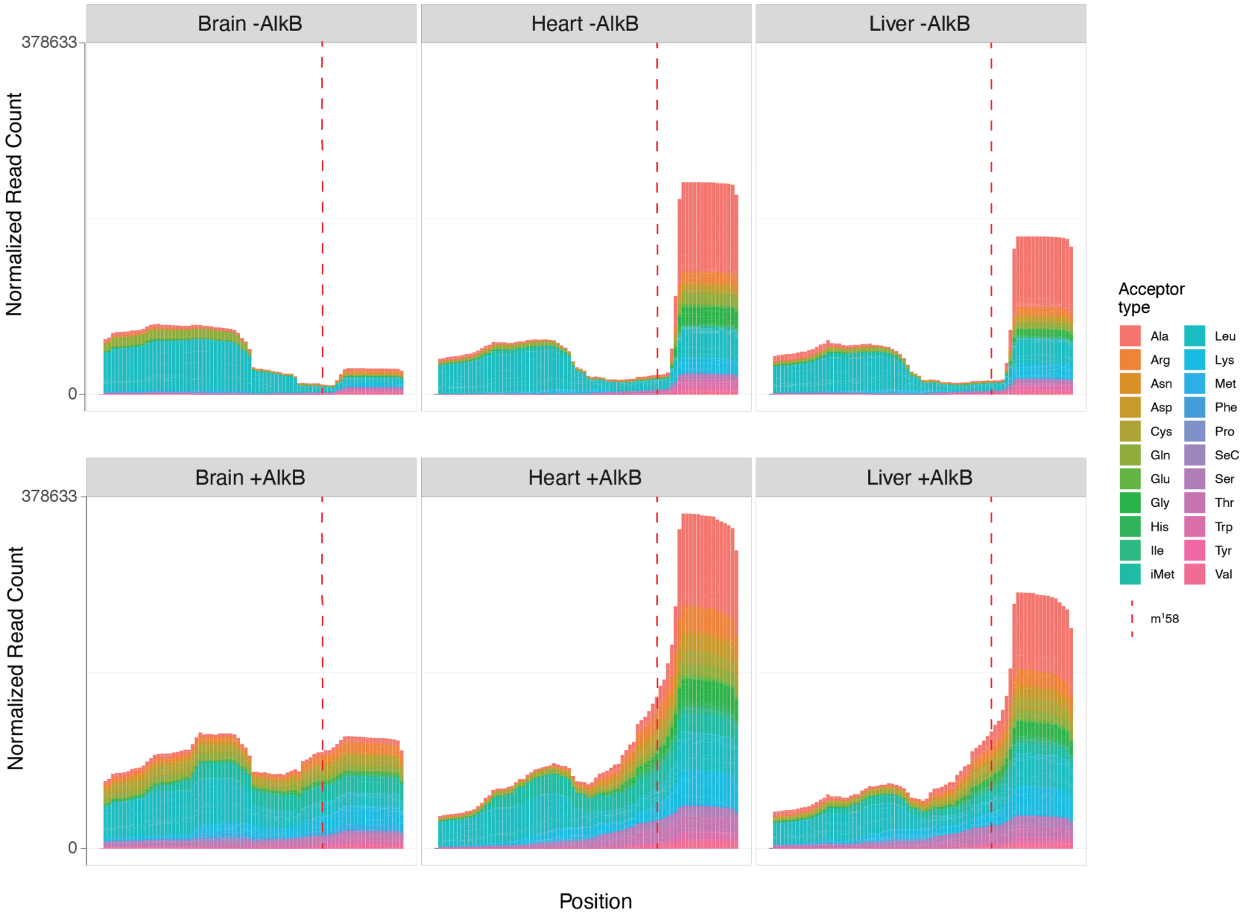


**Figure S9** Combined read coverage across all tRNAs from mouse ARM-seq experiments. Read coverage of all tRNAs categorized by isotype for each tissue type is shown from 5′ end to 3′ end (x-axis). Y-axis represents normalized read count generated by tRAX. Red dotted line represents pos 58 of tRNAs where $m^1A$ modification is commonly found.

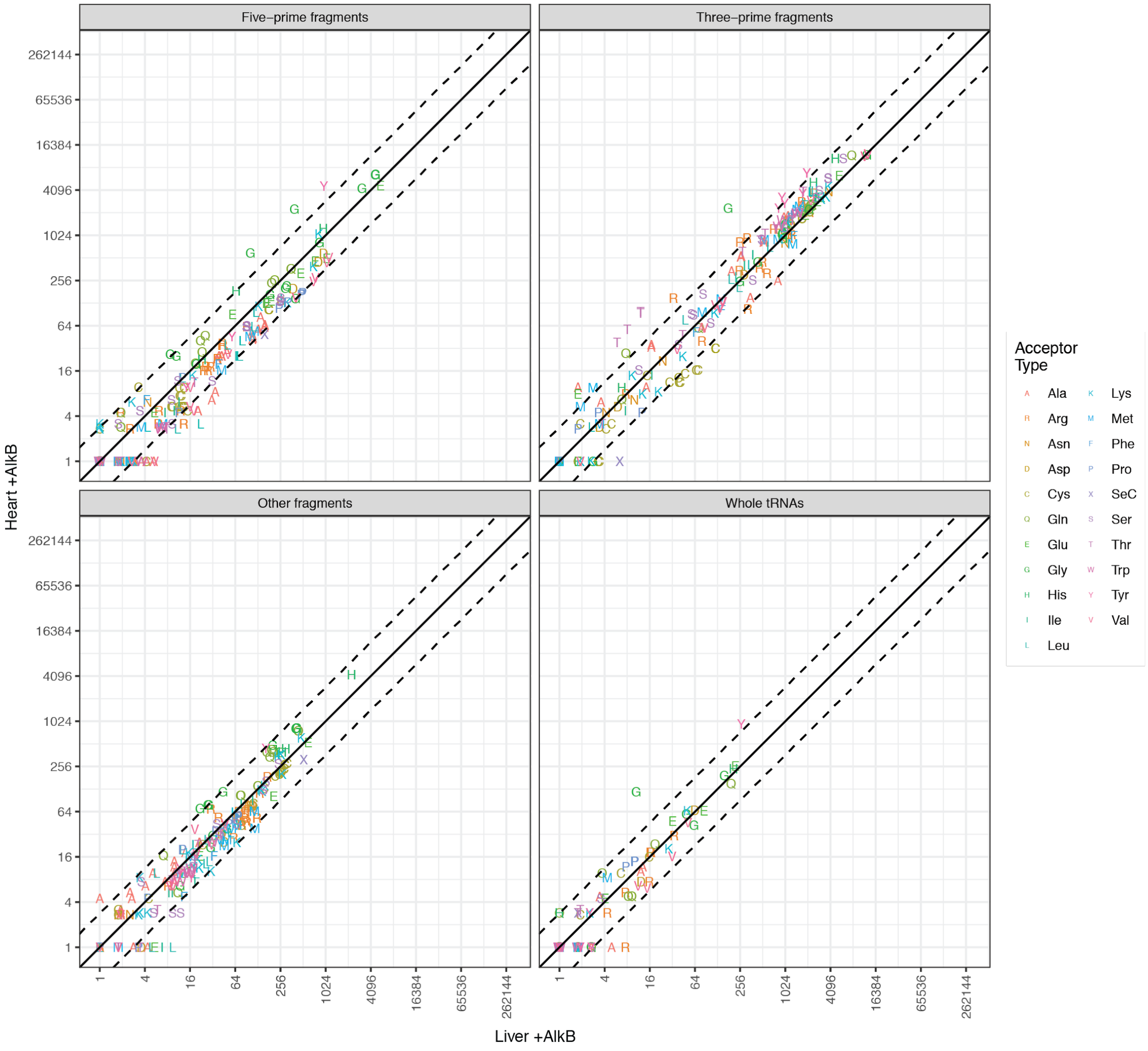


**Figure S10** Mouse heart vs. mouse liver tDR abundance comparison by fragment type. Normalized read counts of AlkB-treated mouse heart and liver samples were used for generating the scatter plots. Counts were categorized into different tRNA fragment types (five-prime fragments, three-prime fragments, and other fragments) to represent expressed tDRs in addition to whole tRNAs. Both tDRs and tRNAs are labeled by tRNA isotype using the amino acid 1-letter code. Solid diagonal lines reflect perfect positive correlation between two samples, while dotted diagonal lines reflect 1.5 $\log_2$-fold differences in abundance between samples.

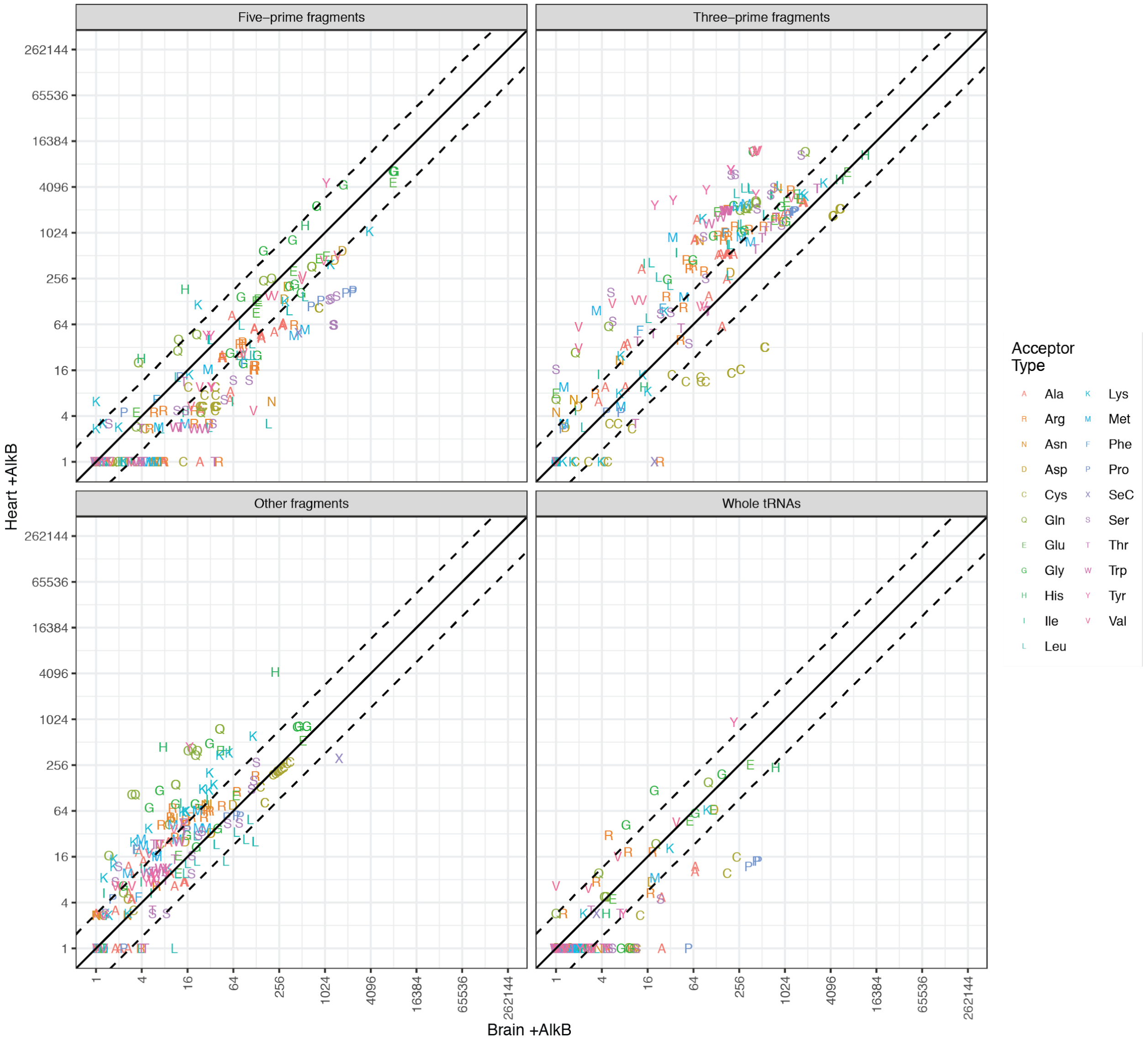


**Figure S11** Mouse heart vs. mouse brain tDR abundance comparison by fragment type. Normalized read counts of AlkB-treated mouse heart and brain samples were used for generating the scatter plots. Counts were categorized into different tRNA fragment types (five-prime fragments, three-prime fragments, and other fragments) to represent expressed tDRs in addition to whole tRNAs. Both tDRs and tRNAs are labeled by tRNA isotype using the amino acid 1-letter code. Solid diagonal lines reflect perfect positive correlation between two samples, while dotted diagonal lines reflect 1.5 $\log_2$-fold differences in abundance between samples.

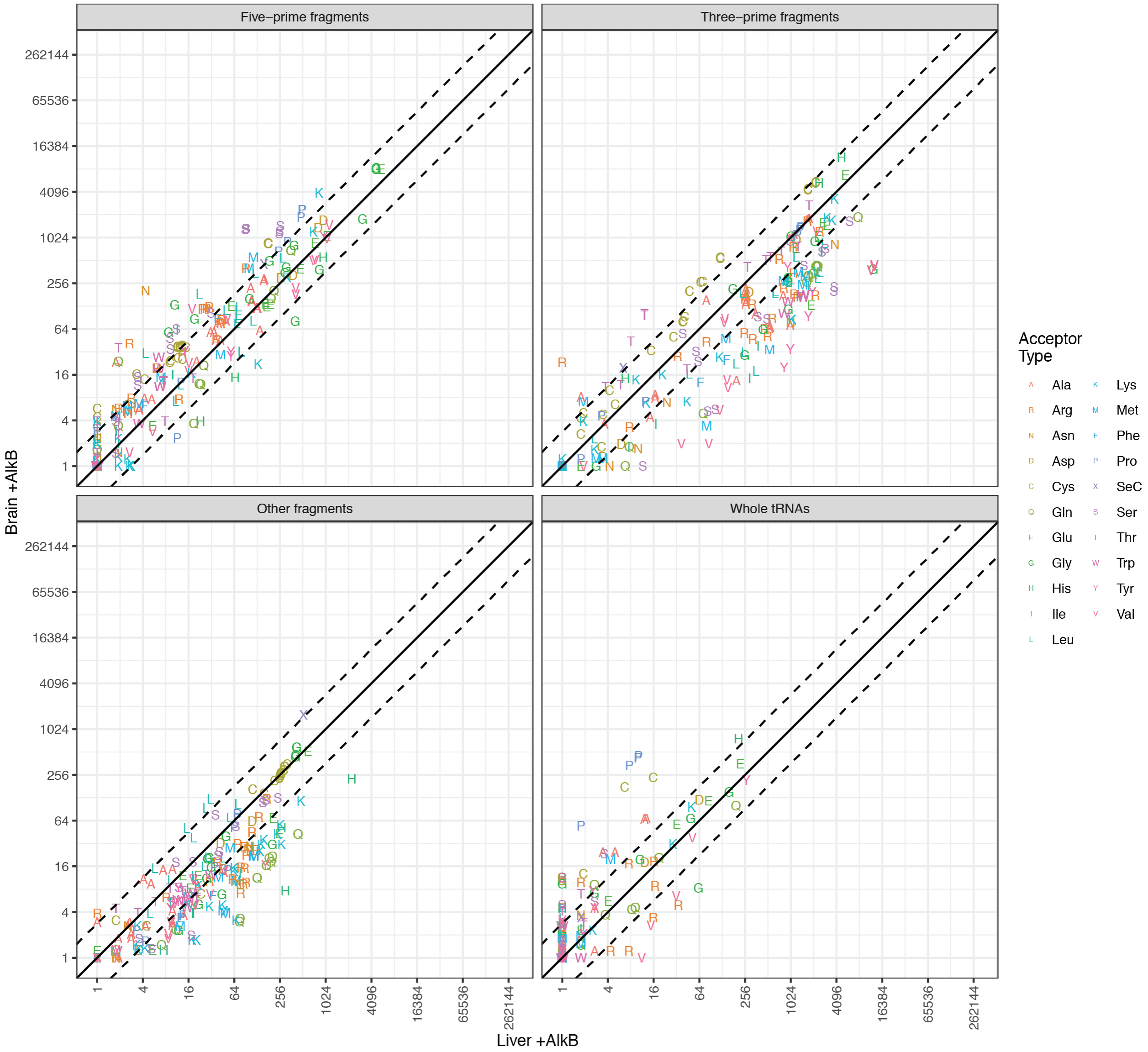


**Figure S12** Mouse brain vs. mouse liver tDR abundance comparison by fragment type. Normalized read counts of AlkB-treated mouse brain and liver samples were used for generating the scatter plots. Counts were categorized into different tRNA fragment types (five-prime fragments, three-prime fragments, and other fragments) to represent expressed tDRs in addition to whole tRNAs. Both tDRs and tRNAs are labeled by tRNA isotype using the amino acid 1-letter code. Solid diagonal lines reflect perfect positive correlation between two samples, while dotted diagonal lines reflect 1.5 $\log_2$-fold differences in abundance between samples.

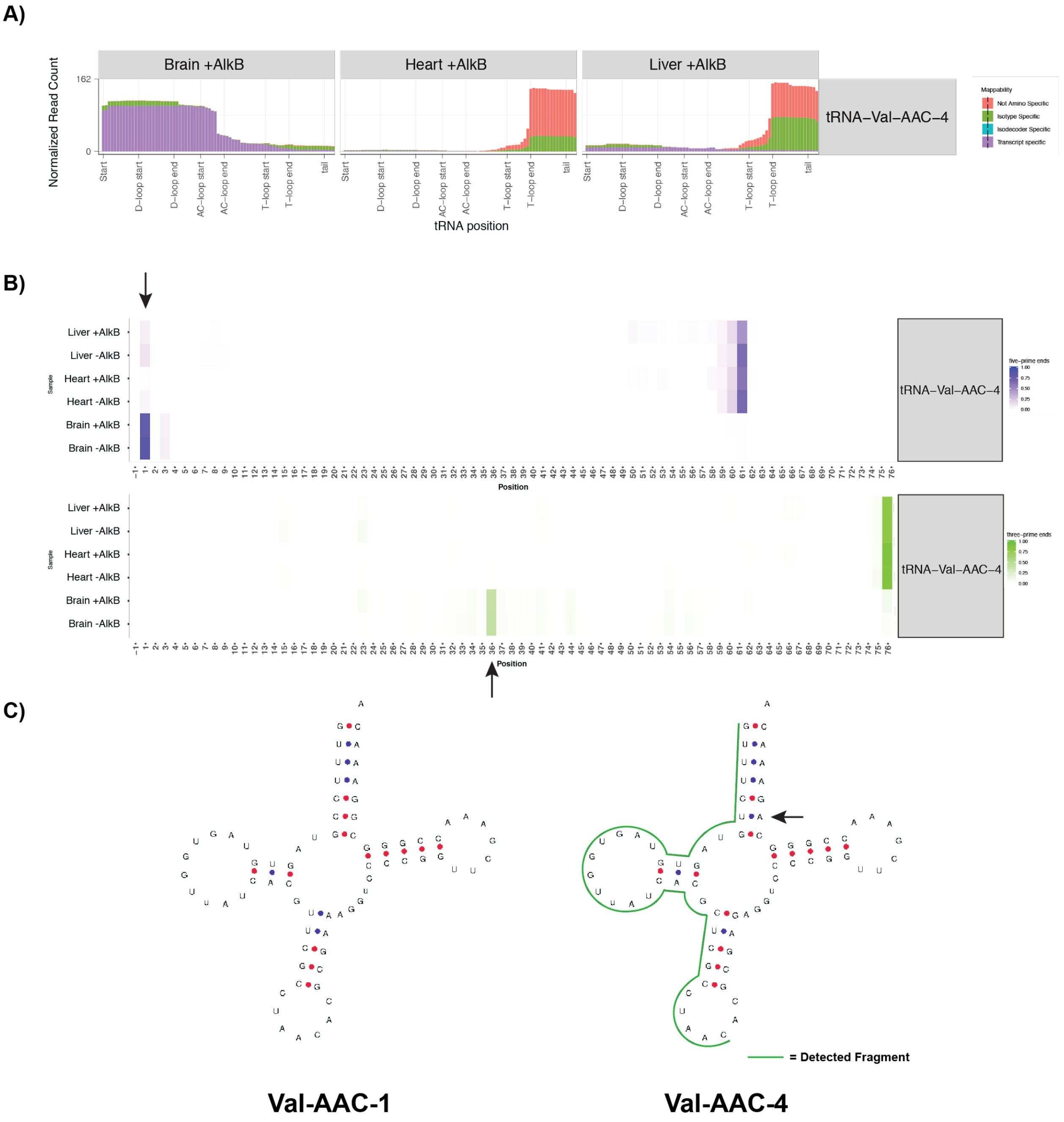


**Figure S13** Brain-enriched tDR derived from tRNA-Val-AAC-4. A) Comparison of read coverage across tRNA-Val-AAC-4 from mouse brain, heart, and liver AlkB-treated samples. B) Heatmaps show the possible 5′ (*top heatmap*) and 3′ (*bottom heatmap*) ends of tDR at positions of tRNA-Val-AAC-4 by the percentage of mapped reads. The 5′ end (position 1) and 3′ end (position 36) of the brain-enriched tDR are indicated with arrows. C) Comparison with tRNA-Val-AAC-1 shows the unique $U_6$:$A_{67}$ at the acceptor stem of tRNA-Val-AAC-4 (marked by arrow). Green line represents the location of the 5′ tDR relative to tRNA-Val-AAC-4.

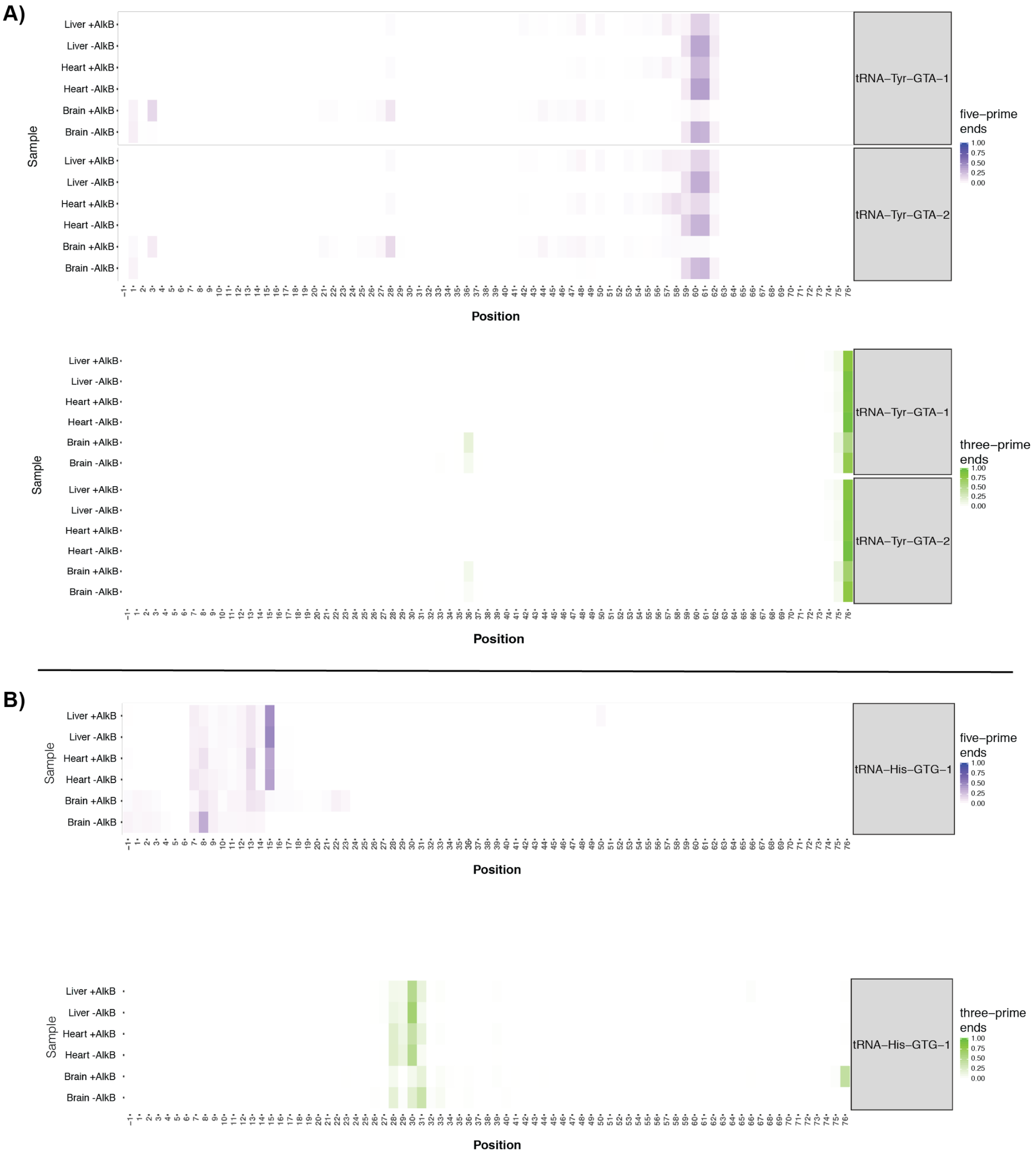


**Figure S14** Mapping of 5′ and 3′ ends of tissue-enriched tDRs derived from tRNA-Tyr-GTA and tRNA-His-GTG. Heatmaps show the possible 5′ (*top heatmap*) and 3′ (*bottom heatmap*) ends of tDRs at positions of A) tRNA-Tyr-GTA isodecoders and B) tRNA-His-GTG-1.

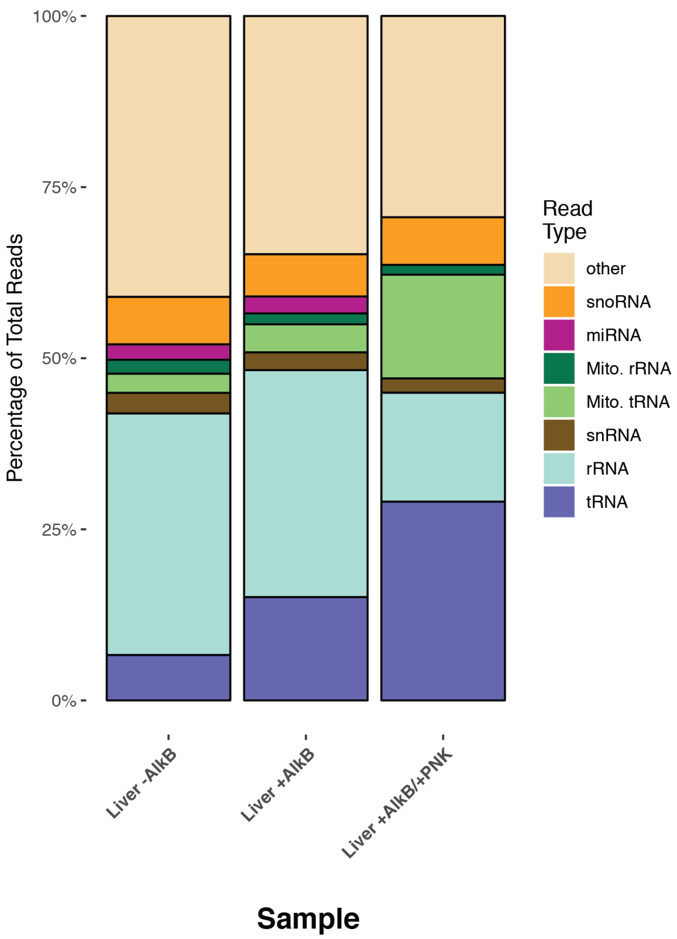


**Figure S15** Sequencing read distribution of mouse liver ARM-seq experiment with T4 polynucleotide kinase treatment (PNK). -AlkB represents untreated samples. +AlkB represents samples with AlkB treatment. +AlkB/+T4PNK represents samples treated with AlkB and T4 PNK.

A)

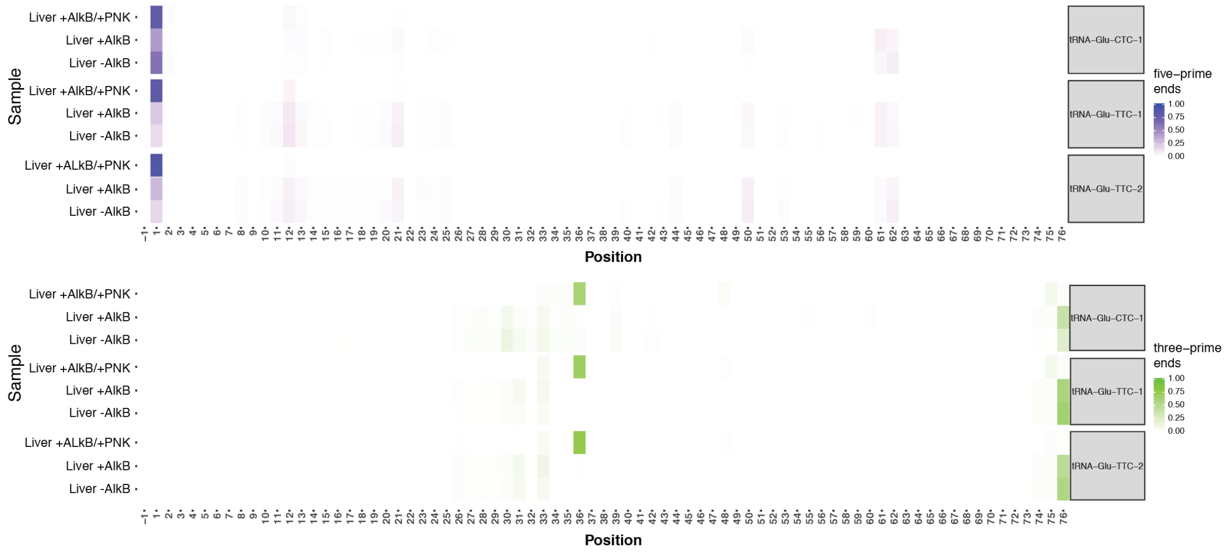


B)

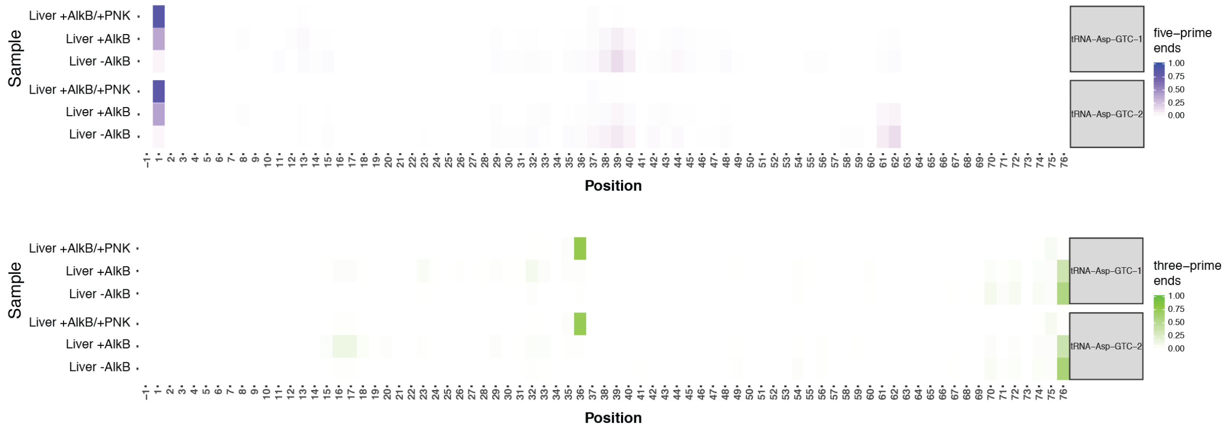


**Figure S16** Mapping of 5′ and 3′ ends of tDRs derived from tRNA-Glu-CTC/TTC and tRNA-Asp-GTC detected in T4 PNK-treated samples. Heatmaps show the possible 5′ (*top heatmap*) and 3′ (*bottom heatmap*) ends of tDRs at positions of A) tRNA-Glu-CTC/TTC isodecoders and B) tRNA-Asp-GTC isodecoders.

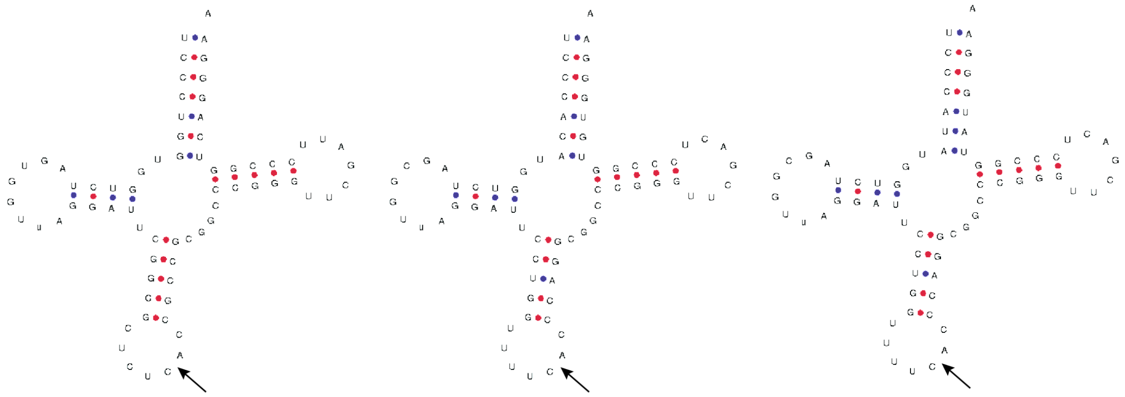

**tRNA-Glu-CTC-1** **tRNA-Glu-TTC-1** **tRNA-Glu-CTC-1**

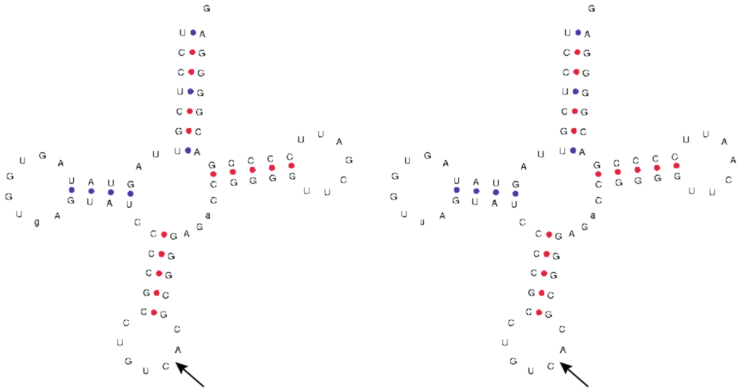

**tRNA-Asp-GTC-1** **tRNA-Asp-GTC-2**

**Figure S17** tDRs detected in T4 PNK-treated samples share the same end position relative to parent tRNAs. Likely endoribonuclease angiogenin cleavage sites are marked by arrows.

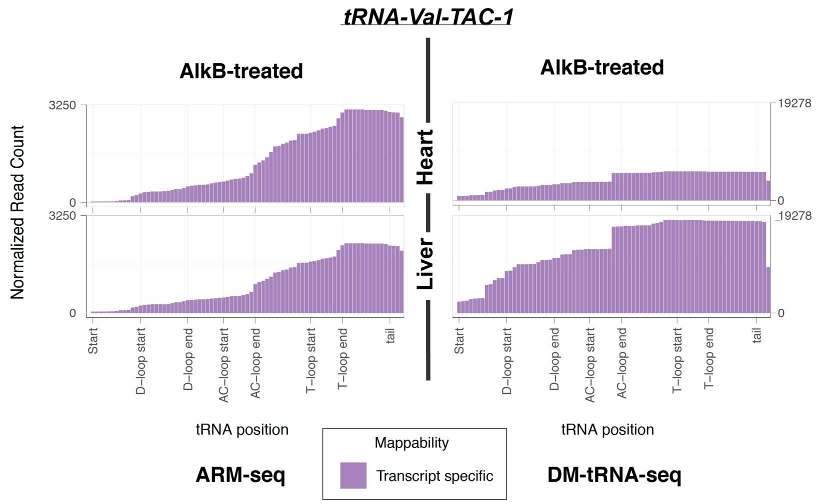


**Figure S18** Read coverage plot comparison for tRNA-Val-TAC-1 between ARM-seq (left) and DM-tRNA-seq (right) in mouse heart (top) and liver (bottom). “Transcript-specific” mappability represents that only sequencing reads uniquely mapped to tRNA-Val-TAC-1 are included in the read coverage.

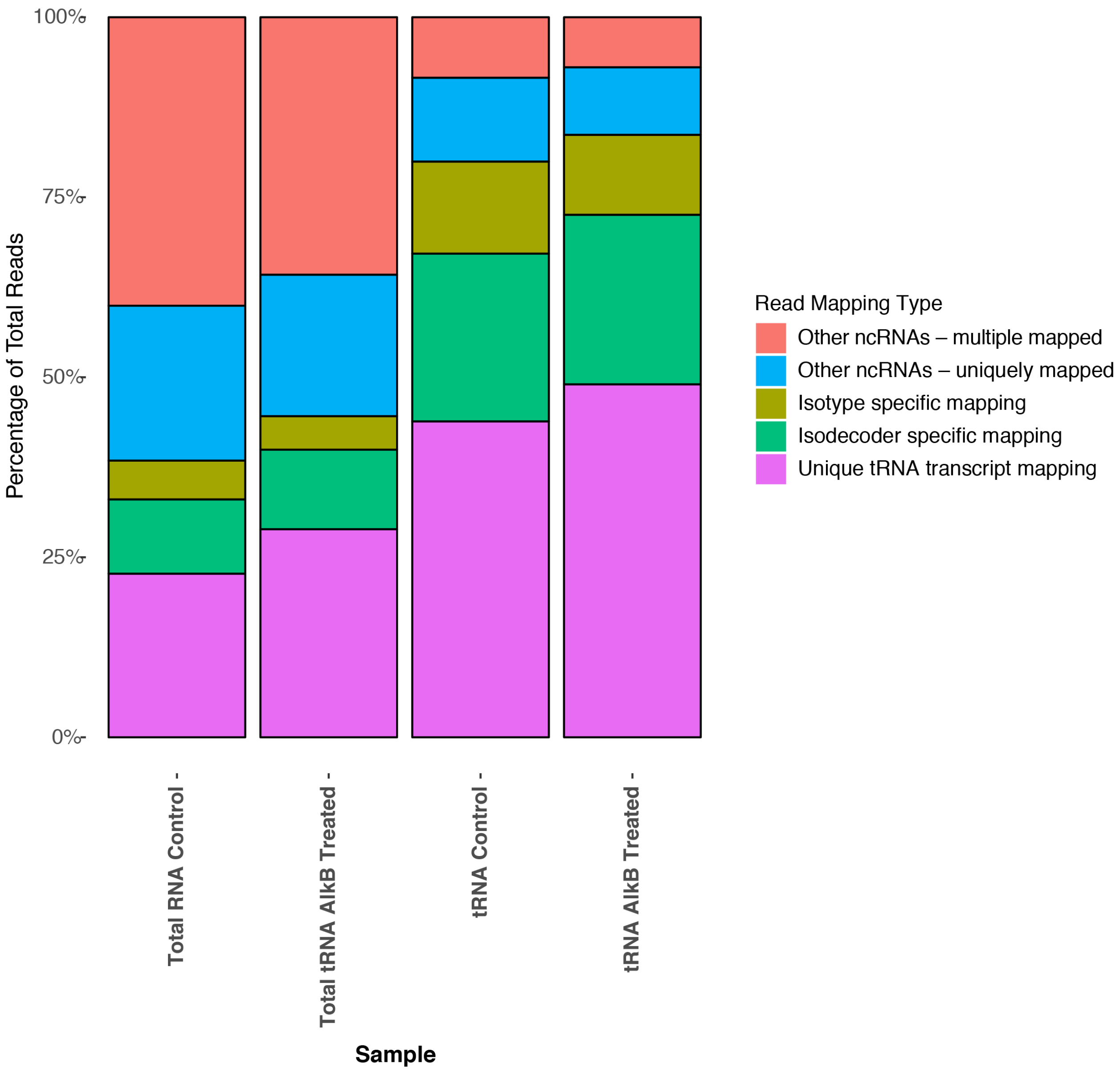


**Figure S19** Read mapping uniqueness distribution of the original DM-tRNA-seq study [1] . Sequencing reads aligned to tRNAs and other ncRNAs were grouped by mappability uniqueness. Reads mapped to multiple tRNA transcripts were further distinguished between isotype-specific (equal matches to tRNAs decoding the same amino acid) and isodecoder-specific (equal matches to multiple different tRNAs with the same anticodon).

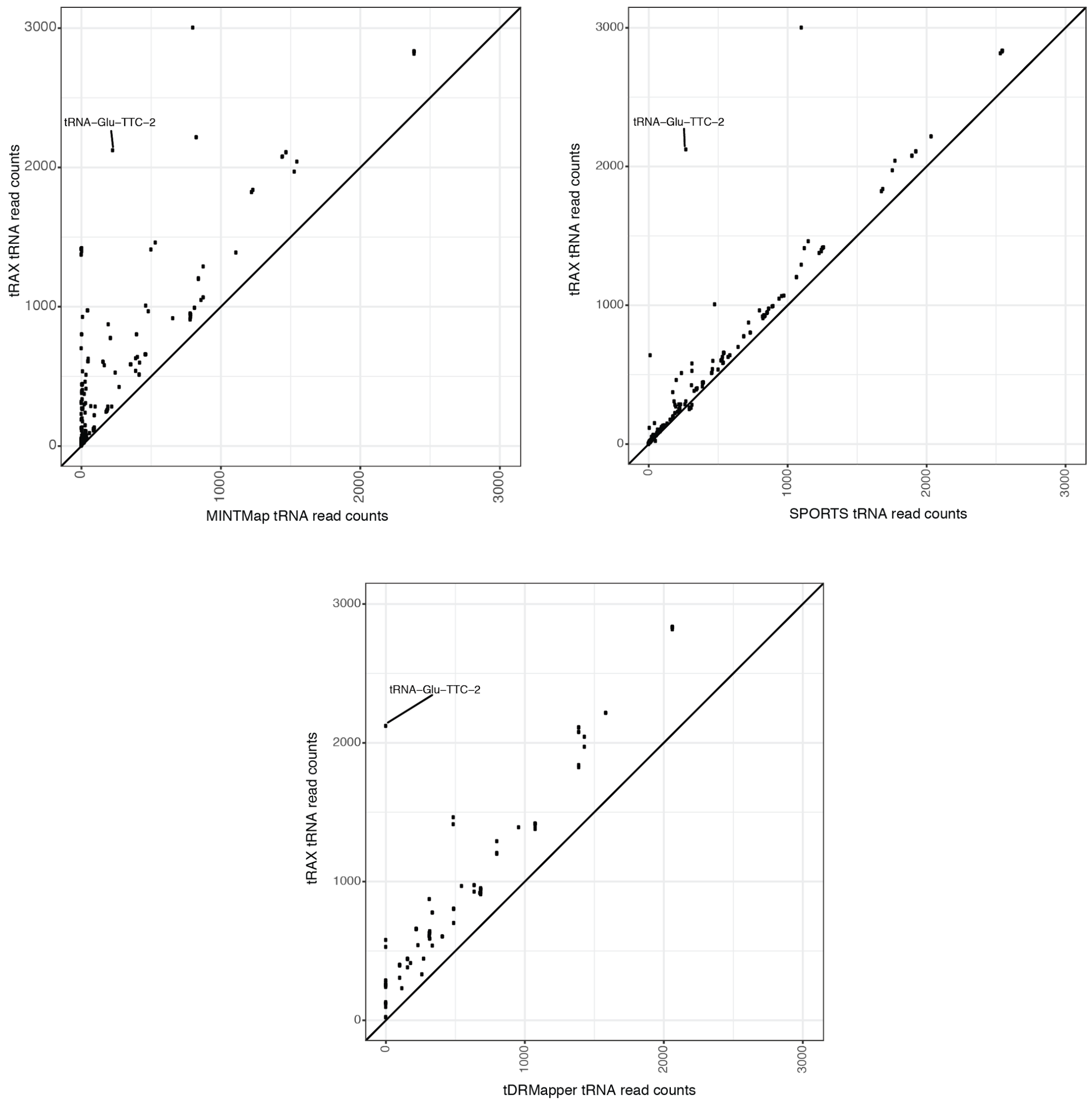


**Figure S20** Read count comparison of tRAX with MINTMap [2], tDRmapper [3], and SPORTS [4] using ARM-seq data. Sequencing data set used in the original ARM-seq study [5] was reanalyzed by tRAX, MINTMap, tDRmapper, and SPORTS. Read counts of tDRs derived from the same mature tRNA were combined for comparison at tRNA level. tRNA-Glu-TTC-2 was highlighted as an example to show the read count difference between tRAX and other tools.

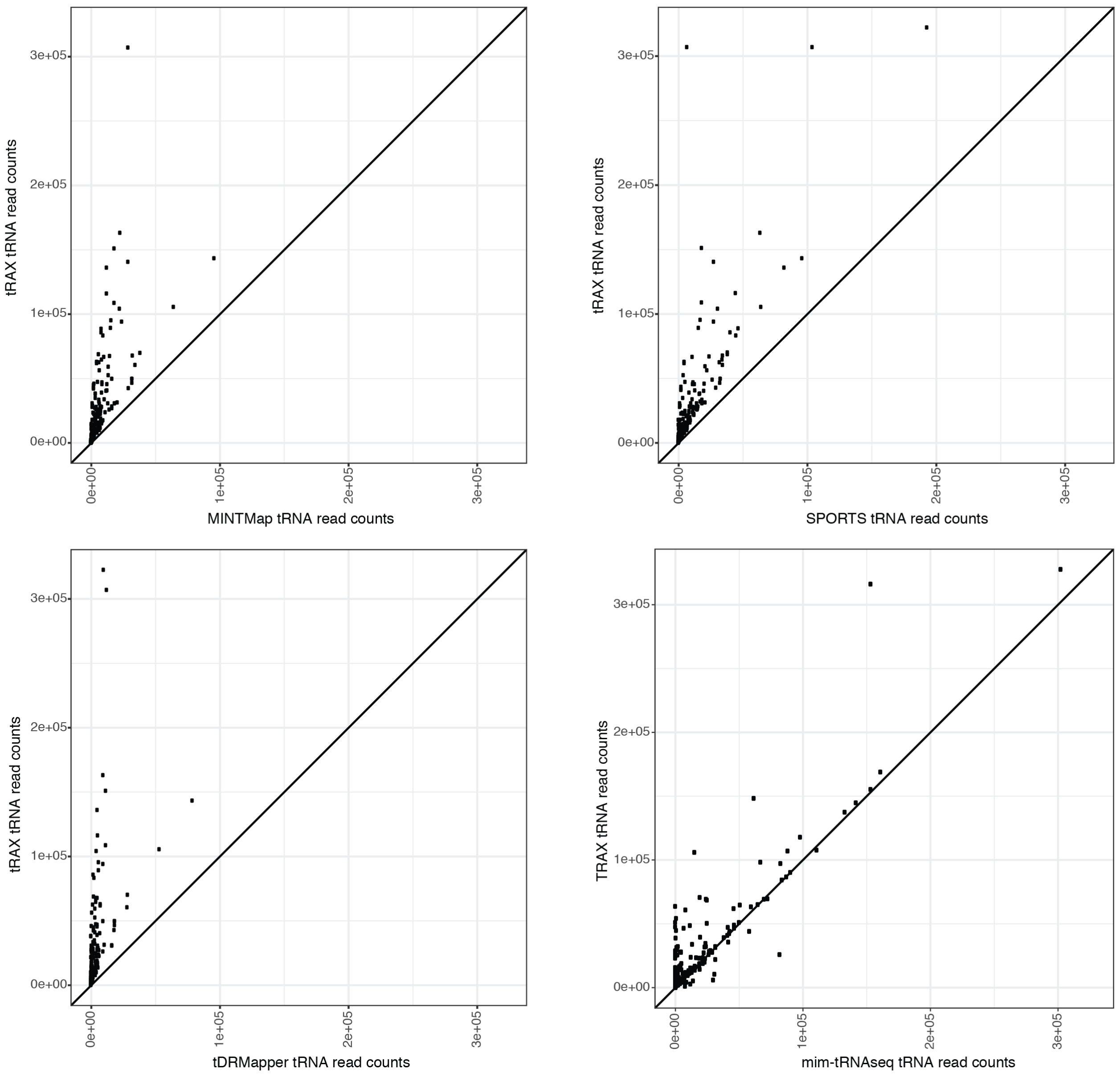


**Figure S21** Read count comparison of tRAX with MINTMap [2], tDRmapper [3], SPORTS [4], and mim-tRNAseq [6] using DM-tRNA-seq data. Sequencing data set used in the original DM-tRNA-seq study [1] was reanalyzed by tRAX, MINTMap, tDRmapper, SPORTS, mim-tRNAseq. Because MINTMap, tDRmapper, and SPORTS only compute read counts for tDRs, all read counts belonging to the same mature tRNA were added together for comparison with tRAX results.
