## Supplementary Data for "tRNA Analysis of eXpression (tRAX): A tool for integrating analysis of tRNAs, tRNA-derived small RNAs, and tRNA modifications"

### tRAX Data Quality Report

**Date: Wednesday September 23, 2020**
**Run mode: Full-length tRNAs**

**Summary**

| Status | Quality Metric | Passed Samples | Warnings | Failures | File |
|---|---|---|---|---|---|
| 12 Warnings | tRNA read share > 50.0% | 6 | 12 | 0 | mousedmtrnaseqreorder-typecounts.pdf |
| Passed | rRNA read share < 35.0% | 18 | 0 | 0 | mousedmtrnaseqreorder-typecounts.pdf |
| Passed | Reads mapping to unannotated regions < 35.0% | 18 | 0 | 0 | mousedmtrnaseqreorder-typecounts.pdf |
| Passed | Read length average < 75 bases | 18 | 0 | 0 | mousedmtrnaseqreorder-readlengths.pdf |
| 15 Warnings | >= 70.0% of reads between 40 and 75 bases | 3 | 15 | 0 | mousedmtrnaseqreorder-readlengths.pdf |
| Passed | >= 50.0% of tRNAs with more than 20 reads | 18 | 0 | 0 | mousedmtrnaseqreorder-tRNAcounts.txt |
| 2 Warnings | DESeq2 size factor differences < 3.0x | 16 | 2 | 0 | mousedmtrnaseqreorder-SizeFactors.txt |

---

**tRNA read share > 50.0%**

| Status | Sample | tRNA Read Percentage |
|---|---|---|
| Passed | M_dm_Liver_M4_plusAlkB | 59.83% |
| Passed | M_dm_Liver_M5_plusAlkB | 55.86% |
| Passed | M_dm_Liver_M6_plusAlkB | 62.04% |
| Passed | M_dm_Liver_M4_minusAlkB | 52.59% |
| Passed | M_dm_Liver_M5_minusAlkB | 53.21% |
| Passed | M_dm_Liver_M6_minusAlkB | 54.60% |
| Warning | M_dm_Heart_M4_plusAlkB | 26.01% |
| Warning | M_dm_Heart_M5_plusAlkB | 32.04% |
| Warning | M_dm_Heart_M6_plusAlkB | 31.90% |
| Warning | M_dm_Heart_M4_minusAlkB | 11.65% |
| Warning | M_dm_Heart_M5_minusAlkB | 16.20% |
| Warning | M_dm_Heart_M6_minusAlkB | 17.36% |
| Warning | Mouse_Brain_M4_plusAlkB | 37.06% |
| Warning | Mouse_Brain_M5_plusAlkB | 42.50% |
| Warning | Mouse_Brain_M6_plusAlkB | 46.19% |
| Warning | Mouse_Brain_M4_minusAlkB | 21.34% |
| Warning | Mouse_Brain_M5_minusAlkB | 25.19% |
| Warning | Mouse_Brain_M6_minusAlkB | 20.61% |

back to Summary

---

**rRNA read share < 35.0%**

| Status | Sample | rRNA Read Percentage |
|---|---|---|
| Passed | M_dm_Liver_M4_plusAlkB | 4.69% |
| Passed | M_dm_Liver_M5_plusAlkB | 4.19% |
| Passed | M_dm_Liver_M6_plusAlkB | 4.52% |
| Passed | M_dm_Liver_M4_minusAlkB | 5.00% |
| Passed | M_dm_Liver_M5_minusAlkB | 3.19% |
| Passed | M_dm_Liver_M6_minusAlkB | 3.59% |
| Passed | M_dm_Heart_M4_plusAlkB | 3.26% |
| Passed | M_dm_Heart_M5_plusAlkB | 2.50% |
| Passed | M_dm_Heart_M6_plusAlkB | 2.51% |
| Passed | M_dm_Heart_M4_minusAlkB | 2.21% |
| Passed | M_dm_Heart_M5_minusAlkB | 1.54% |
| Passed | M_dm_Heart_M6_minusAlkB | 1.58% |
| Passed | Mouse_Brain_M4_plusAlkB | 2.25% |
| Passed | Mouse_Brain_M5_plusAlkB | 2.45% |
| Passed | Mouse_Brain_M6_plusAlkB | 3.51% |
| Passed | Mouse_Brain_M4_minusAlkB | 4.32% |
| Passed | Mouse_Brain_M5_minusAlkB | 3.80% |
| Passed | Mouse_Brain_M6_minusAlkB | 3.90% |

back to Summary

---

**Reads mapping to unannotated regions < 35.0%**

| Status | Sample | Unannotated Region Mapping Rate |
|---|---|---|
| Passed | M_dm_Liver_M4_plusAlkB | 21.88% |
| Passed | M_dm_Liver_M5_plusAlkB | 28.10% |
| Passed | M_dm_Liver_M6_plusAlkB | 22.98% |
| Passed | M_dm_Liver_M4_minusAlkB | 19.11% |
| Passed | M_dm_Liver_M5_minusAlkB | 21.62% |
| Passed | M_dm_Liver_M6_minusAlkB | 17.99% |
| Passed | M_dm_Heart_M4_plusAlkB | 27.51% |
| Passed | M_dm_Heart_M5_plusAlkB | 26.54% |
| Passed | M_dm_Heart_M6_plusAlkB | 23.39% |
| Passed | M_dm_Heart_M4_minusAlkB | 19.27% |
| Passed | M_dm_Heart_M5_minusAlkB | 19.00% |
| Passed | M_dm_Heart_M6_minusAlkB | 17.24% |
| Passed | Mouse_Brain_M4_plusAlkB | 22.04% |
| Passed | Mouse_Brain_M5_plusAlkB | 19.43% |
| Passed | Mouse_Brain_M6_plusAlkB | 19.69% |
| Passed | Mouse_Brain_M4_minusAlkB | 20.97% |
| Passed | Mouse_Brain_M5_minusAlkB | 19.85% |
| Passed | Mouse_Brain_M6_minusAlkB | 21.49% |

back to Summary

---

**Read length average < 75 bases**

| Status | Sample | Average Read Length |
|---|---|---|
| Passed | M_dm_Liver_M4_plusAlkB | 52.65 |
| Passed | M_dm_Liver_M5_plusAlkB | 59.32 |
| Passed | M_dm_Liver_M6_plusAlkB | 61.76 |
| Passed | M_dm_Liver_M4_minusAlkB | 59.89 |
| Passed | M_dm_Liver_M5_minusAlkB | 62.18 |
| Passed | M_dm_Liver_M6_minusAlkB | 60.88 |
| Passed | M_dm_Heart_M4_plusAlkB | 66.31 |
| Passed | M_dm_Heart_M5_plusAlkB | 63.92 |
| Passed | M_dm_Heart_M6_plusAlkB | 53.95 |
| Passed | M_dm_Heart_M4_minusAlkB | 61.07 |
| Passed | M_dm_Heart_M5_minusAlkB | 60.42 |
| Passed | M_dm_Heart_M6_minusAlkB | 58.00 |
| Passed | Mouse_Brain_M4_plusAlkB | 60.91 |
| Passed | Mouse_Brain_M5_plusAlkB | 62.89 |
| Passed | Mouse_Brain_M6_plusAlkB | 60.77 |
| Passed | Mouse_Brain_M4_minusAlkB | 56.79 |
| Passed | Mouse_Brain_M5_minusAlkB | 58.27 |
| Passed | Mouse_Brain_M6_minusAlkB | 60.78 |

back to Summary

---

**>= 70.0% of reads between 40 and 75 bases**

| Status | Sample | Read Percentage |
|---|---|---|
| Warning | M_dm_Liver_M4_plusAlkB | 52.47% |
| Warning | M_dm_Liver_M5_plusAlkB | 60.56% |
| Warning | M_dm_Liver_M6_plusAlkB | 57.59% |
| Warning | M_dm_Liver_M4_minusAlkB | 61.28% |
| Warning | M_dm_Liver_M5_minusAlkB | 67.93% |
| Warning | M_dm_Liver_M6_minusAlkB | 66.36% |
| Warning | M_dm_Heart_M4_plusAlkB | 61.18% |
| Warning | M_dm_Heart_M5_plusAlkB | 54.33% |
| Warning | M_dm_Heart_M6_plusAlkB | 53.37% |
| Warning | M_dm_Heart_M4_minusAlkB | 62.61% |
| Warning | M_dm_Heart_M5_minusAlkB | 60.30% |
| Warning | M_dm_Heart_M6_minusAlkB | 59.38% |
| Passed | Mouse_Brain_M4_plusAlkB | 70.26% |
| Warning | Mouse_Brain_M5_plusAlkB | 67.05% |
| Warning | Mouse_Brain_M6_plusAlkB | 68.30% |
| Warning | Mouse_Brain_M4_minusAlkB | 66.82% |
| Passed | Mouse_Brain_M5_minusAlkB | 70.39% |
| Passed | Mouse_Brain_M6_minusAlkB | 70.59% |

back to Summary

---

**>= 50.0% of tRNAs with more than 20 reads**

| Status | Sample | Percentage of tRNAs |
|---|---|---|
| Passed | M_dm_Liver_M4_plusAlkB | 96.79% |
| Passed | M_dm_Liver_M5_plusAlkB | 96.79% |
| Passed | M_dm_Liver_M6_plusAlkB | 96.79% |
| Passed | M_dm_Liver_M4_minusAlkB | 96.79% |
| Passed | M_dm_Liver_M5_minusAlkB | 96.79% |
| Passed | M_dm_Liver_M6_minusAlkB | 96.79% |
| Passed | M_dm_Heart_M4_plusAlkB | 96.79% |
| Passed | M_dm_Heart_M5_plusAlkB | 96.79% |
| Passed | M_dm_Heart_M6_plusAlkB | 96.79% |
| Passed | M_dm_Heart_M4_minusAlkB | 96.79% |
| Passed | M_dm_Heart_M5_minusAlkB | 96.79% |
| Passed | M_dm_Heart_M6_minusAlkB | 96.79% |
| Passed | Mouse_Brain_M4_plusAlkB | 96.79% |
| Passed | Mouse_Brain_M5_plusAlkB | 96.79% |
| Passed | Mouse_Brain_M6_plusAlkB | 96.79% |
| Passed | Mouse_Brain_M4_minusAlkB | 96.79% |
| Passed | Mouse_Brain_M5_minusAlkB | 96.79% |
| Passed | Mouse_Brain_M6_minusAlkB | 96.79% |

back to Summary

---

**DESeq2 size factor differences < 3.0x**

| Status | Sample | Size Factor |
|---|---|---|
| Warning | M_dm_Liver_M4_plusAlkB | 0.13 |
| Passed | M_dm_Liver_M5_plusAlkB | 1.91 |
| Passed | M_dm_Liver_M6_plusAlkB | 1.85 |
| Passed | M_dm_Liver_M4_minusAlkB | 1.33 |
| Passed | M_dm_Liver_M5_minusAlkB | 0.80 |
| Passed | M_dm_Liver_M6_minusAlkB | 1.30 |
| Passed | M_dm_Heart_M4_plusAlkB | 2.19 |
| Passed | M_dm_Heart_M5_plusAlkB | 0.66 |
| Passed | M_dm_Heart_M6_plusAlkB | 1.61 |
| Passed | M_dm_Heart_M4_minusAlkB | 1.04 |
| Passed | M_dm_Heart_M5_minusAlkB | 2.03 |
| Passed | M_dm_Heart_M6_minusAlkB | 1.63 |
| Passed | Mouse_Brain_M4_plusAlkB | 1.58 |
| Passed | Mouse_Brain_M5_plusAlkB | 2.10 |
| Passed | Mouse_Brain_M6_plusAlkB | 1.35 |
| Passed | Mouse_Brain_M4_minusAlkB | 0.55 |
| Passed | Mouse_Brain_M5_minusAlkB | 0.96 |
| Warning | Mouse_Brain_M6_minusAlkB | 0.10 |

back to Summary

---
